# Lead-DBS v2: Towards a comprehensive pipeline for deep brain stimulation imaging

**DOI:** 10.1101/322008

**Authors:** Andreas Horn, Ningfei Li, Till A Dembek, Ari Kappel, Chadwick Boulay, Siobhan Ewert, Anna Tietze, Andreas Husch, Thushara Perera, Wolf-Julian Neumann, Marco Reisert, Hang Si, Robert Oostenveld, Christopher Rorden, Fang-Cheng Yeh, Qianqian Fang, Todd M Herrington, Johannes Vorwerk, Andrea A. Kühn

**Author notes:** Authors contributed equally (Andreas Horn, Ningfei Li). Corresponding Author Dr. Andreas Horn Movement Disorders & Neuromodulation Unit Department for Neurology Charité – University Medicine Berlin Charitéplatz 1, 10117 Berlin, Germany.

## Abstract

Deep brain stimulation (DBS) is a highly efficacious treatment option for movement disorders and a growing number of other indications are investigated in clinical trials. To ensure optimal treatment outcome, exact electrode placement is required. Moreover, to analyze the relationship between electrode location and clinical results, a precise reconstruction of electrode placement is required, posing specific challenges to the field of neuroimaging. Since 2014 the open source toolbox Lead-DBS is available, which aims at facilitating this process. The tool has since become a popular platform for DBS imaging. With support of a broad community of researchers worldwide, methods have been continuously updated and complemented by new tools for tasks such as multispectral nonlinear registration, structural / functional connectivity analyses, brain shift correction, reconstruction of microelectrode recordings and orientation detection of segmented DBS leads. The rapid development and emergence of these methods in DBS data analysis require us to revisit and revise the pipelines introduced in the original methods publication. Here we demonstrate the updated DBS and connectome pipelines of Lead-DBS using a single patient example with state-of-the-art high-field imaging as well as a retrospective cohort of patients scanned in a typical clinical setting at 1.5T. Imaging data of the 3T example patient is co-registered using five algorithms and nonlinearly warped into template space using ten approaches for comparative purposes. After reconstruction of DBS electrodes (which is possible using three methods and a specific refinement tool), the volume of tissue activated is calculated for two DBS settings using four distinct models and various parameters. Finally, four whole-brain tractography algorithms are applied to the patient’s preoperative diffusion MRI data and structural as well as functional connectivity between the stimulation volume and other brain areas are estimated using a total of eight approaches and datasets. In addition, we demonstrate impact of selected preprocessing strategies on the retrospective sample of 51 PD patients. We compare the amount of variance in clinical improvement that can be explained by the computer model depending on the method of choice.

This work represents a multi-institutional collaborative effort to develop a comprehensive, open source pipeline for DBS imaging and connectomics, which has already empowered several studies, and may facilitate a variety of future studies in the field.

## Introduction

In the field of deep brain stimulation (DBS), precise electrode placement is crucial for optimal treatment outcomes. Specifically, a direct relationship between electrode localization and clinical outcome has been shown in multiple studies (e.g. Butson et al., 2011; Dembek et al., 2017; Eisenstein et al., 2014; Garcia-Garcia et al., 2016; Horn et al., 2017c; Mosley et al., 2018b; also see fig. 1 A). To characterize this relationship in an objective manner, tools are required that facilitate the reconstruction of electrode placement such that comparisons between patients can be made. Group comparisons play a crucial role in identifying optimal electrode placement, providing both direct clinical and theoretical insights. Ideally, to fulfill reproducibility and transparency criteria needed for good scientific practice, these tools should be open source and publicly available. Finally, a specific challenge that differentiates the field of DBS imaging from most other neuroimaging domains is the need for absolute anatomical precision. A shift of two mm in electrode placement may represent a major change in clinical outcome, while in conventional fMRI studies, a change of an activity peak by two mm has little if no impact at all (figure 1).

**Figure 1:**
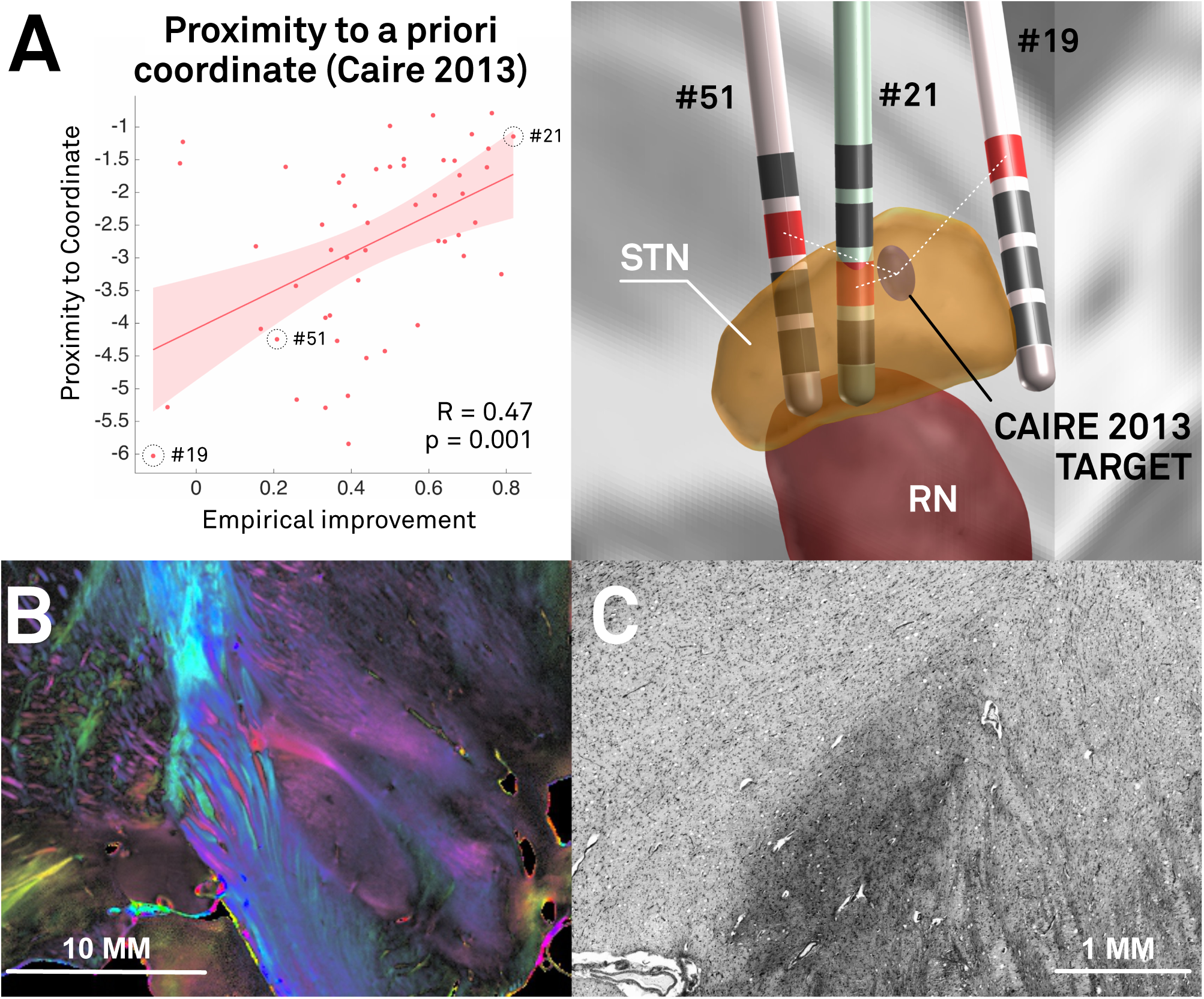
In DBS imaging, “millimeters matter”, which poses specific methodological challenges. A) Re-analysis of the Berlin cohort described in (Horn et al., 2017c) shows that proximity of active contact centers to an optimal target is predictive of %-UPDRS-III improvement (left). The target was defined in a meta-analysis (Caire et al., 2013) and transformed to MNI space in a probabilistic fashion (Horn et al., 2017a). Active contacts between electrodes of patients 51 and 21 are a mere two mm away from each other, but result in largely different clinical results (right). The same distance (two mm) corresponds to the average image resolution of functional MRI for which many neuroimaging tools were initially developed. Thus, in the field of DBS imaging, the distance of two mm plays a crucial role, whereas it is often considered insignificant in common neuroimaging studies. B) Coronal polarized light imaging section of the human subthalamic nucleus with surrounding tracts. Image courtesy by Prof. Karl Zilles and Dr. Markus Axer, Forschungszentrum Jülich, INM-1. C) Coronal section of the BigBrain dataset (Amunts et al., 2013) as visualized in the microdraw online application (http://microdraw.pasteur.fr/). Cell sparser and denser subregions are discernible, potentially corresponding to functional zones of the nucleus (Marani et al., 2008). B) and C) demonstrate the tightly-packed anatomical complexity of the STN-DBS target region that is similarly reflected in clinical outcome (A). Of note, only a subregion of this small nucleus is considered an optimal DBS target. The combination of such small and complex DBS targets with a potentially huge impact of small misplacements poses extreme challenges to the field of DBS imaging and raise the need for high-precision pipelines.

In 2014, the software toolbox Lead-DBS was published that aimed at reconstructing DBS electrode placement based on pre- and postoperative imaging (Horn and Kühn, 2015; www.lead-dbs.org; RRID:SCR_002915). Using the toolbox, electrodes may be localized in relationship to surrounding brain anatomy. Since its initial publication, development efforts have continued at multiple institutions. Thus, over the years, numerous progress has been made and better alternatives for most steps described in the original pipeline are now provided (Ewert et al., 2018a; 2018b; Horn et al., 2017a; 2017b; 2017c). Moreover, several novel features that were not mentioned (or available) in the original publication have recently become crucial components of DBS imaging. These have now been integrated in the latest release. While other tools with similar aims have been introduced after publication of Lead-DBS (Bonmassar et al., 2014; da Silva et al., 2015; D’Albis et al., 2014; Husch et al., 2017; 2018; Lauro et al., 2015), the tool was recently described as the most established toolbox for electrode localizations (Husch et al., 2017) with over 6600 downloads and 75 citations. The aim of the project is to develop a scientific platform in a multi-institutional endeavor that is and remains available under an open license (GNU general public license v. 3) to ensure reproducibility and version control.

The growing user base of Lead-DBS as an academic toolbox and the divergence of the current methods and those described in the initial publication raise the need of an updated methodological pipeline description. In addition, we also use the opportunity to emphasize the latest default analysis options, pitfalls and methods throughout the pipeline.

Given the complexity of multiple processing stages (see figure 2 & tables 1-4), a thorough empirical evaluation of each stage exceeds the scope of this work. For instance, it would represent a study in itself to empirically probe which normalization method, which stimulation volume model or which fiber tracking approach could yield best results. Such studies have been conducted (Åström et al., 2014; Dembek et al., 2017; Fillard et al., 2011; Klein et al., 2009; Maier-Hein et al., 2017; McIntyre et al., 2004) and are currently underway in context of the Lead-DBS environment, as well (Ewert et al., 2018a). Instead, the aim of the present article is to give an overview of methods available in Lead-DBS. To make the processing stages concrete, the pipeline is described using a single patient example with state-of-the art high-field (3T) imaging as well as a retrospective sample of 51 PD patients imaged at 1.5T. The result is a focus on the methods section and a descriptive results section covering co-registration, normalization, electrode localization, VTA estimation, and structural-functional connectivity analyses. Finally, we demonstrate that more variance in clinical outcome may be explained when using the default pipeline in comparison to a more “standard neuroimaging” approach. The manuscript has a narrative prose with the aim of maximizing understandability while omitting unnecessary details where possible. Moreover, while the manuscript is still structured into conventional sections, the methods descriptions exceed the actual processing of the study with the aim of illustrating the multiple approaches implemented in Lead-DBS and providing notes about motivation and potential limitations.

**Figure 2:**
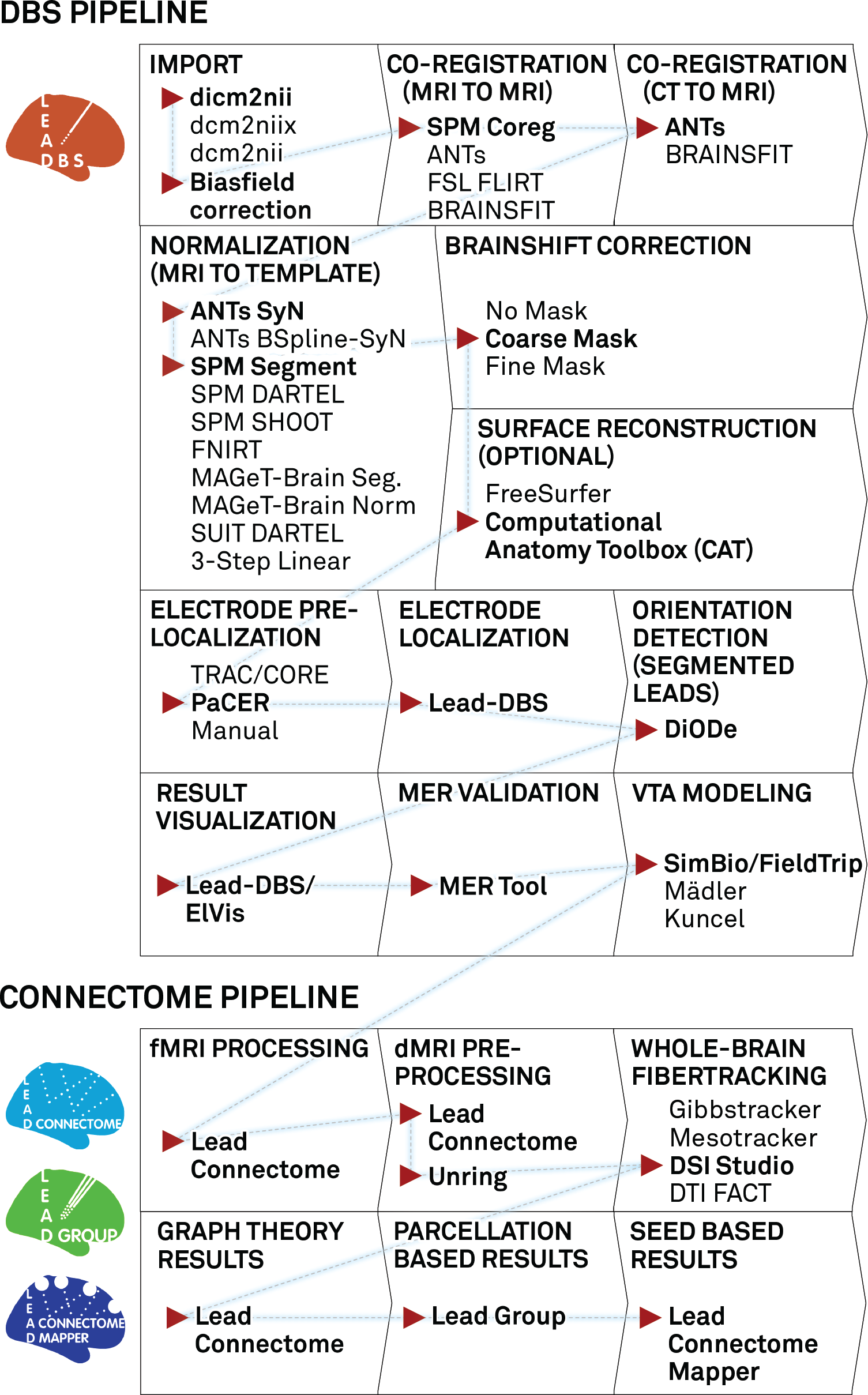
A “default pathway” through Lead-DBS. Processing stages are visualized in consecutive order, general choices are displayed for each step while default selections are marked with red arrow and bold text. For the normalization step, a larger evaluation showed both the ANTs SyN and SPM Segment approaches to perform equally optimal (Ewert et al., 2018a). The DBS and connectome pipelines work in parallel but seamlessly integrate via the order marked by the blue dashed line. After calculating results with Lead Connectome Mapper (last box), results may be used to predict clinical outcome using the Lead Predict tool (not shown) based on a model described in (Horn et al., 2017c).

**Table 1:**
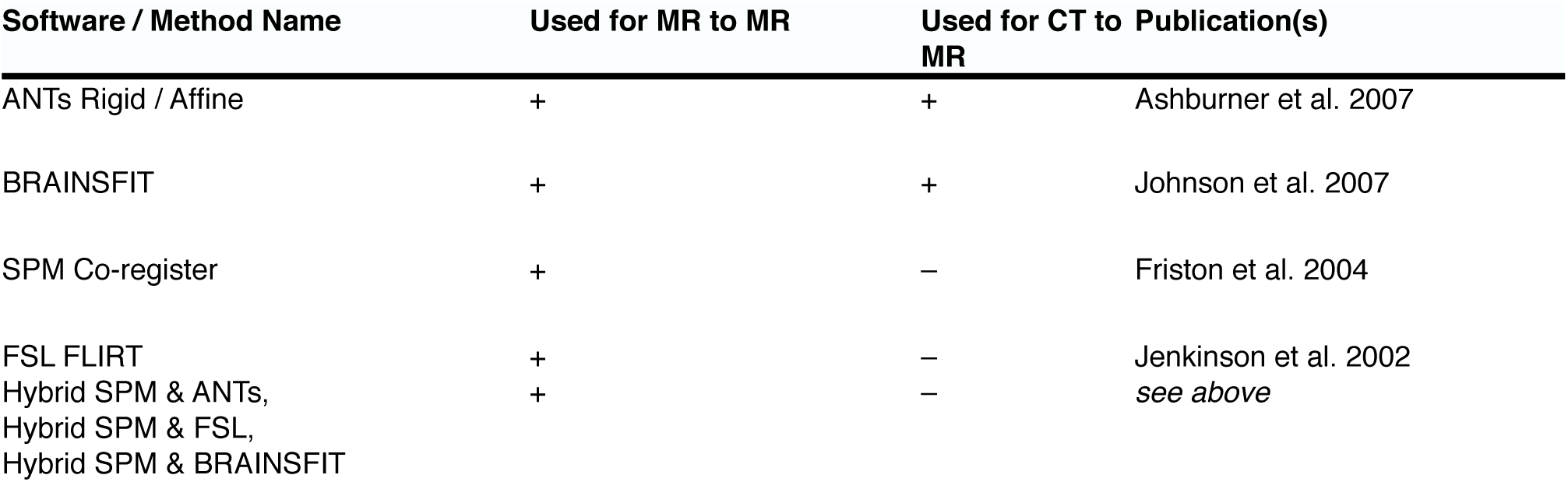
Linear registration methods implemented in Lead-DBS.

| Software / Method Name | Used for MR to MR | Used for CT to MR | Publication(s) |
| --- | --- | --- | --- |
| ANTs Rigid / Affine | + | + | Ashburner et al. 2007 |
| BRAINSFIT | + | + | Johnson et al. 2007 |
| SPM Co-register | + | – | Friston et al. 2004 |
| FSL FLIRT | + | – | Jenkinson et al. 2002 |
| Hybrid SPM & ANTs,<br>Hybrid SPM & FSL,<br>Hybrid SPM & BRAINSFIT | + | – | *see above* |

## Methods

### Patient Characteristics, Surgery & Imaging Example patient

A male patient (65y) suffering from Parkinson‘s Disease received two octopolar segmented DBS leads (Boston Scientific Vercise; BSci, Marlborough, Massachusetts, United States) targeting the subthalamic nucleus (STN). Surgery was done under general anaesthesia with two wakeful phases in which microelectrode recordings were obtained using a Neuro Omega drive (Alpha Omega Engineering, Nazareth, Israel) with a 45° rotated Ben Gun array. In the same session, test stimulations were performed. Recordings started 7.5 mm before reaching the target and were acquired in 15 consecutive steps of 0.5 mm. Recordings of cells with typical STN firing patterns were later transferred to the Lead-DBS session. Test stimulations were made at three mm dorsal to and at the surgical target. In a second surgery five days afterwards, a Boston Scientific Vercise Gevia Impulse Generator was implanted in the chest. Detailed imaging parameters can be found in supplementary material.

### Retrospective patient cohort

Data from the patient described above embodies a state-of-the-art example dataset acquired at 3T including a specialized basal ganglia MR sequence and patient-specific diffusion MRI. To further illustrate the impact that different processing streams may have on typical clinical MRI data, we included data from a priorly published retrospective cohort that is described in detail elsewhere (Berlin cohort in Horn et al., 2017c). In brief, 51 patients received quadropolar electrodes (Medtronic type 3389) to the STN region to treat Parkinson‘s Disease. Pre- and postoperative imaging was performed on a 1.5T MRI and included a preoperative T1 and T2 sequence as well as postoperative axial, coronal and sagittal T2 slabs of the basal ganglia. Six of 51 patients received a postoperative CT instead (for detailed imaging parameters see supplementary material and Horn et al., 2017c).

### Linear (Within-Patient) Co-Registrations

When a patient folder is loaded in Lead-DBS, a bias-field correction step based on the N4 algorithm is automatically applied to all pre-operative MRI sequences (Tustison et al., 2010). Based on configuration preferences, Lead-DBS chooses one of the preoperative sequences as the anchor modality, i.e. the stationary sequence to which all other (preoperative and postoperative volumes) sequences are co-registered. By default, the T1-weighted sequence is used or if unavailable the T2-weighted sequence is substituted. This anchor modality is upsampled to isotropic 0.7 mm resolution to maintain high resolution in following steps. This step is common in similar pipelines (e.g. Gunalan et al., 2017). A reason is that (e.g. T2 weighted) acquisitions acquired in clinical routine often come in high in-plane resolution (e.g. 0.5 mm) but poor slice thickness (2-3 mm). If these images are resliced to a 1 mm isotropic MP-RAGE, much in-plane resolution is lost. Thus, our pipeline compromises on a 0.7 mm isotropic working space to which the anchor modality is resliced (images need to be resliced for multispectral normalizations). Several linear registration algorithms are included in Lead-DBS (see table 1). In the present example patient and retrospective cohort, all available preoperative acquisitions (i.e. T2, PD, FGATIR as well as the FA volume derived from the dMRI scan) were co-registered and resliced to the upsampled T1 using SPM 12 (https://www.fil.ion.ucl.ac.uk/spm/software/spm12/). Similarly, the postoperative CT was co-registered using Advanced Normalization Tools (ANTs; http://stnava.github.io/ANTs/). Co-registration results were then manually checked using built-in tools that facilitate visual inspection which may be enhanced by automatic edge detection based wire-frame generation of the anchor modality, and false-color overlays.

Co-registration is a crucial step to achieve precise results since the preoperative data is used to define anatomy and the postoperative to define electrode locations. Thus, imprecise registrations lead to erroneous results in the end. In clinical settings, especially on MRI, postoperative volumes are often slabs (i.e. don‘t cover the whole brain). Accurately registering these to preoperative data is especially challenging for registration algorithms and at times, postoperative data must be registered manually (e.g. using tools as 3D Slicer; www.slicer.org).

### Nonlinear (Patient-to-Template) Co-Registrations

To relate electrode placement to anatomy and to make them comparable across patients and centers, it is useful to register individual patient anatomy to a template space. These template spaces often allow the most likely location of anatomical structures to be better defined, and can then be used to project subcortical atlases (table 5) or whole-brain parcellations (table 6) onto regions of interest.

In the original Lead-DBS publication, all registrations between patient and template space were performed in a linear way following the approach introduced by Schönecker and colleagues (Schönecker et al., 2009). This approach was especially developed for DBS and followed a three-step registration with incremental focus on the subcortical region of interest. In the revised version of Lead-DBS, multiple nonlinear options have been added (table 2). Most of the included approaches were adapted with their parameters tuned for optimal results in the DBS context. Most refinements were performed based on experience in the daily use of the pipeline across multiple institutions. Recently, a study systematically analyzed results of various methods in which over 11,000 nonlinear deformations were solved and compared (Ewert et al., 2018a). Results of this study led to the present default presets implemented in Lead-DBS (figure 2). It is beyond the scope of the present work to describe every modification in each method in detail but the ability to explain clinical improvement using some examples was estimated for the retrospective cohort analyzed here and some details about modifications in regard to subcortical optimization are mentioned in the supplementary material.

**Table 2:** Normalization Methods implemented in Lead-DBS:

| Software / Method Name | Adaptations in Lead-DBS | Supports<br>Multispectral<br>Warps | Publication(s) |
| --- | --- | --- | --- |
| **Statistic Parametric Mapping (SPM)** |  |  |  |
| Unified Segmentation | Based on Tissue Probability Maps calculated from multispectral ICBM 2009b NLIN ASYM Space templates | + | Ashburner and Friston 2005 |
| DARTEL | Pair-wise instead of group-wise workflow (mapping directly from patient into template space) based on similarly generated DARTEL templates | + | Ashburner et al. 2007 |
| SHOOT | See DARTEL | + | Ashburner and Friston 2011 |
| SUIT based DARTEL | Specialized method if area of interest is in the brainstem (e.g. PPN); Use of SUIT Toolbox based registration to a brainstem/cerebellum template that was registered to ICBM 2009b | + | Diedrichsen et al. 2006, Ashburner et al. 2007 |
| **FMRIB Software Library (FSL)** |  |  |  |
| FNIRT | Standard presets | – | Andersson and Smith 2010 |
| **Advanced Normalization Tools (ANTs)** |  |  |  |
| SyN | Four-Stage preset with subcortical refinement, multiple presets developed for use in DBS | + | Avants et al. 2008 |
| BSpline-SyN | See SyN | + | Tustison & Avants 2013 |
| **Other / Specialized** |  |  |  |
| Linear Three-Step Normalization | Suited for use directly on postoperative MRI (or on preoperative MRI); Implemented using ANTs whereas original code was implemented with use of FSL | + | Schonecker et al. 2009 |
| MAGeT Brain-like Segmentation / Normalization | Multi-Subject/Template implementation inspired by the MAGeT-Brain approach but with strong differences. | + | None / see Chakravarty et al. 2012 for MAGeT-Brain approach |
| MAGeT Brain-like Normalization | Inversing the idea of MAGeT-Brain by directly averaging deformation fields learned from multiple indirect warps | + | None / see Chakravarty et al. 2012 for MAGeT-Brain approach |

### Brain Shift Correction

During surgery, air may enter the skull after it is opened. This leads to a nonlinear deformation of the brain in relation to the bone which is called brain shift and typically pushes the forebrain into occipital direction (due to supine position of the patient). Especially when the pneumocephalus is still present during postoperative imaging, it introduces a bias between electrode (postoperative image) placement and anatomical structures on preoperative acquisitions. Whilst brain-shift introduces non-linear transforms, applying non-linear registration techniques to correct this would also deform the electrodes projections and corrupt the corresponding anatomical overlap. To avoid this, our brain-shift correction method uses the threefold linear registration described above (see Schönecker et al., 2009 for validation), which is stored internally and applied to DBS electrode placement afterwards (figure S1).

### Electrode Localization

In Lead-DBS, the process of reconstructing electrode placement is divided into an automated (“pre-localization”) and a manual (“localization”) step. A wide range of electrodes from five manufacturers are readily implemented in Lead-DBS (table 7) and it is straight forward to implement custom models.

#### Automated pre-Localization

For the pre-localization part, four methods are available:

1. Manual click-and-point tool
2. Integration with 3D Slicer in which fiducial points are placed manually.
3. TRAC / CORE approach (Horn and Kühn, 2015)
4. The PaCER toolbox (https://adhusch.github.io/PaCER/; Husch et al., 2017).

In practice, PaCER usually requires very little to no manual refinement compared to the TRAC/CORE algorithm, but requires a postoperative CT acquisition in contrast to the other methods.

#### Manual Localization

The user-interface of this crucial processing step was specifically designed to allow for highly precise electrode reconstructions. At all times, the postoperative volume is visualized along planes that are cut orthogonally to the electrode (when moving the lead reconstruction in space, these cuts are updated in real time). A specialized “x-ray mode” can be activated in which the same view is enhanced by averaged stacks of orthogonal slices surrounding the lead. This visualization mode is helpful to reconstruct electrodes in poor resolution acquisitions where partial volume effects may blur or even shift the electrode artifact in space.

### Estimating the Local Volume of Tissue Activated

The Volume of Tissue Activated (VTA) is a conceptual volume that is thought to elicit additional action potentials due to the electrical stimulations of axons (McIntyre and Grill, 2002). Much work has been done in this regard and models with increasing sophistication were introduced over the years (e.g. Åström et al., 2014; Butson and McIntyre, 2008; Chaturvedi et al., 2013). In contrast, some more clinically oriented papers aimed at finding fast heuristics to determine the rough extent of the VTA based on the stimulation parameters without actually creating a spatial model (Dembek et al., 2017; Kuncel et al., 2008; Lauro et al., 2015; Mädler and Coenen, 2012). Three such simple heuristic models are included in Lead-DBS (Dembek et al., 2017; Kuncel et al., 2008; Mädler and Coenen, 2012), and a more sophisticated, finite element method based approach was added in 2017 (Horn et al., 2017c; table 3). On a spectrum between simple heuristical and highly sophisticated models, it falls in the middle. This model was briefly described in the aforementioned publication and follows the overall concept described in (Åström et al., 2014). However, a full methods description of the specific model has not been published and can be found in the supplementary material.

**Table 3:** VTA models included in Lead-DBS.

| Method Publication | Type |
| --- | --- |
| Horn et al. 2017 | FEM Based, 4-Compartment Model, Tetrahedral Mesh |
| Mädler & Coenen 2012 | Heuristic |
| Kuncel et al. 2008 | Heuristic |
| Dembek et al. 2017 | Heuristic |

### Connectivity Estimation

In the revised version of Lead-DBS, several possibilities to estimate structural and functional connectivity exist. These are accessible via the submodules Lead Connectome, Lead Group and Lead Connectome Mapper. Moreover, single-patient connectivity metrics may be visually explored in the general 3D viewer module of the software (ElVis).

Methods may be divided into approaches that utilize *patient-specific* vs. *normative/group-level* resting-state functional MRI (rs-fMRI) or diffusion-weighted imaging based tractography (dMRI). Moreover, they can be subdivided in *voxel-wise / data-driven* or *parcellation-/ROI-based* methods.

#### Patient Specific Connectivity Estimates

A structural-functional processing pipeline was implemented into the Lead Connectome submodule based on the pipeline described in (Horn, 2015; Horn and Blankenburg, 2016; Horn et al., 2014). For rs-fMRI, the pipeline follows the recommendations given in (Weissenbacher et al., 2009). Briefly, this includes motion-correction (SPM based), detrending, regression of white matter and CSF-signals as well as motion parameters and bandpass-filtering (cutoff-values: 0.009-0.08 Hz) of time series. In dMRI, a Gibbs’ ringing removal step (Kellner et al., 2015) is performed before passing the diffusion data into either of four tools subsequently used to estimate a whole-brain tractogram (for a list of fiber-tracking methods implemented in Lead-DBS, see table 4). The Gibbs’ tracking algorithm was superior to nine competing algorithms in the 2009 Fiber Cup (Fillard et al., 2011) and was added as the first method. Its successor, a model-free version (Konopleva et al., 2018) of the Mesotracker algorithm (Reisert et al., 2014) was subsequently added to equally investigate mesoscopic properties of fiber tracts and support multi-shell diffusion data. Recently, the Generalized q-Sampling Imaging approach (Yeh et al., 2010) implemented in DSI Studio (http://dsi-studio.labsolver.org/) achieved the highest “valid connection” score in an open competition among 96 methods submitted from 20 different research groups around the world (Maier-Hein et al., 2017). As a result, this method was included into Lead-DBS as well. Finally, a dated simple tensor based deterministic method is also available for debugging or testing purposes.

**Table 4:**
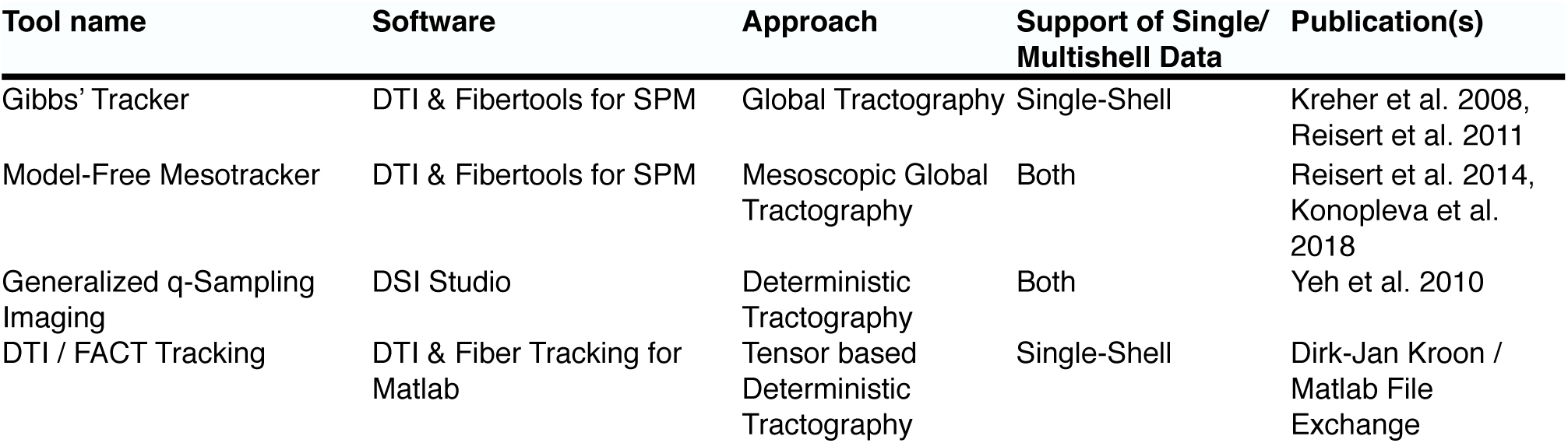
Whole-Brain fiber tracking methods implemented in Lead-DBS. Of note, the tensor based method is implemented for debugging purposes and not recommended for actual use.

**Table 5:**
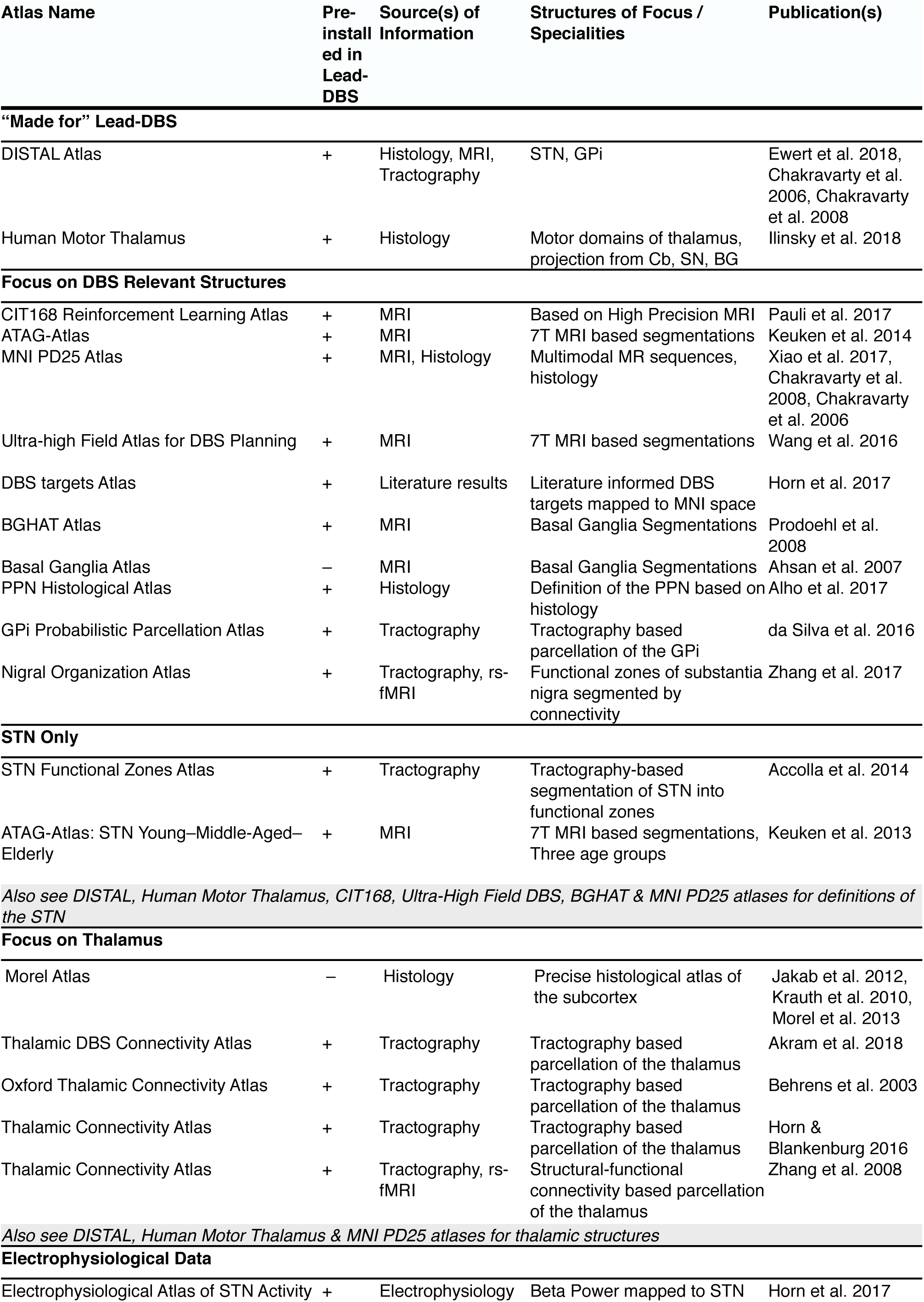
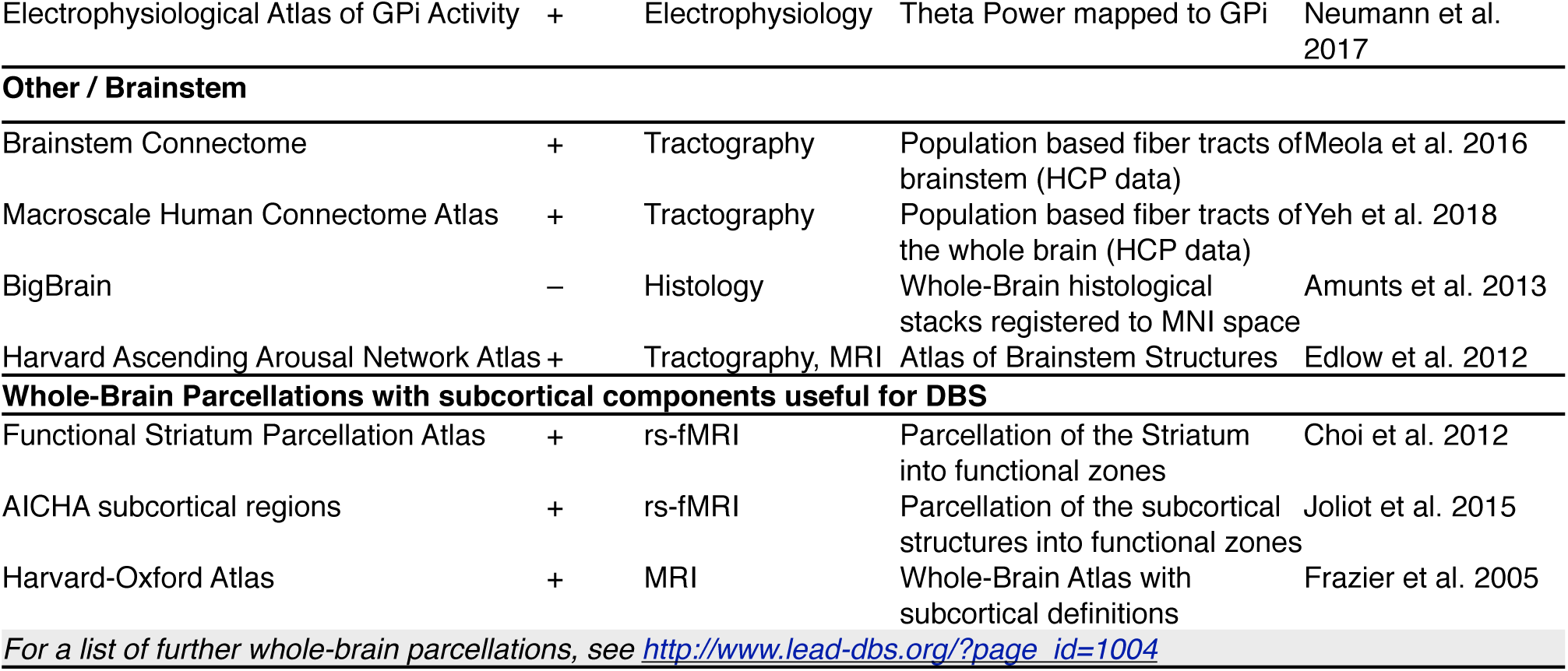
Subcortical Atlases suitable for / available within Lead-DBS.

#### Normative or population based Connectomes

In the DBS field, large cohorts exist in which patient-specific connectivity data is lacking. In such datasets, a novel technique that combines normative group connectome data with single-patient imaging results may be used. These group connectomes were informed by large cohorts of subjects or patients (e.g. N = 1000 in case of the Yeo 2011 normative connectome) that were often acquired on specialized MR hardware (such as the human connectome scanner at the Athinoula A. Martinos Center, Boston, MA; Setsompop et al., 2013). The utility of such normative connectomes in a clinical context was first demonstrated by mapping various neurological or psychiatric symptoms to networks influenced by stroke lesions (Boes et al., 2015; Darby et al., 2016; 2017; Fasano et al., 2017; Fischer et al., 2016; Laganiere et al., 2016). Recently, the approach was adapted to the field of DBS in first studies (Horn et al., 2017c; 2017b; 2017a) and, in order, to predict clinical outcome of transcranial magnetic stimulation treatment (Weigand et al., 2017).

A natural limitation of the approach is that the normative connectome data does not account for patient-specific differences in brain connectivity. However, despite its potential shortcomings in individualized connectivity, the use of normative connectomes has a major practical advantage, since larger cohorts of DBS patients with individualized connectivity data are not available and connectivity sequences in DBS patients are difficult to acquire postoperatively. As such, this approach is able to utilize large DBS cohorts collected across different centers, while studies using patient-specific connectivity are based on small cohorts (typically N < 25; e.g. Accolla et al., 2016; Akram et al., 2017; Vanegas Arroyave et al., 2016). Along the same lines, the approach may prove particularly valuable for emerging DBS indications in which only a limited number of patients are implanted world-wide. Thus, the ability to retrospectively analyze such DBS datasets despite the lack of patient-specific connectivity data represents a genuine window into understanding the role of brain connectivity in mediating DBS outcome.

The methods utilizing normative connectomes are implemented in the Lead Connectome Mapper, Lead Group and ElVis tools (figure 2). For an overview of normative connectomes available within Lead-DBS, see table 8.

### Explaining variance in clinical outcome within the retrospective 1.5T cohort

DBS-Electrodes of the retrospective cohort were localized using Lead-DBS (see fig. 2) after several normalization and registration strategies were performed. Specifically, preoperative acquisitions were registered into template space using the two default approaches (SPM New Segment and ANTs SyN) identified in (Ewert et al. 2018a). In addition, the ANTs SyN approach without the subcortical refinement step was applied. Finally, a T1-only monospectral approach (FSL FNIRT) was added to compare results with a more typical “standard procedure” used in the neuroimaging field. Electrode localizations (performed in native space) where then registered to template space using the deformation fields obtained by the various approaches and VTAs were calculated using default parameters in template space. Overlaps between VTAs and the subthalamic nucleus as defined by the DISTAL atlas (Ewert et al. 2018b) were calculated for both hemispheres and summed up. In addition, overlap between the E-field and the STN were calculated in a weighted fashion by multiplying the binary STN image with the non-binary E-Field and summing up voxels. Finally, streamlines were isolated from a priorly published normative connectome based on the Parkinson‘s Progression Marker Inititative (PPMI) data (Marek et al. 2011; Ewert et al. 2018) using the E-Field of the SPM New Segment method as a weighted seed.

These imaging based metrics (VTA-STN overlap, E-Field-STN weighted overlap, weighted streamlines seeding from E-Field connected to SMA) were correlated with empirical % improvement on the Unified Parkinson‘s Disease Rating Scale (UPDRS) III. In a second step, they were fed into general linear models (GLM) that additionally included seven additional clinical covariates of the sample (age, sex, percent improvement in Levodopa response, disease duration until surgery, Levodopa Equivalent Daily Dosage (LEDD) ON and OFF DBS as well as %LEDD reduction by DBS). From these GLMs, root-mean-square error (RSME), R^2^-Statistic and p-value of the F-statistic as well as significance predictors are reported.

### Methods summary

A list of tools to which interfaces exist or that form a native (preinstalled) part of Lead-DBS is given in table 10. To make deliberate choices regarding which option to choose for each processing step, users require a high methodological level of understanding. To account for this, figure 2 gives an overview of the “default pathway” through Lead-DBS which is further demonstrated in detail in a walkthrough-video available online (http://www.lead-dbs.org/?page_id=192).

## Results

### Patient Outcome

#### 3T example patient

Before surgery, the 64 year male patient had an UPDRS-III score of 64 points (OFF dopamine replacement therapy; Hoehn-Yahr stage IV). Seven days post-surgery, the patient was discharged with an appreciable stun effect and a subjective improvement of gait. Stimulation was set to 0.5 mA bilateral on the lower segmented contacts (ring mode). Under stimulation and medication, a UPDRS-III of four points was taken. At the time of writing, no score under dopaminergic withdrawal and under stimulation is available but will be taken during the three-month postoperative visit.

#### Retrospective 1.5T cohort

The 51 (18 female) patients were 60 ± 7.9 years old at time of surgery and UPRDS-III improved by 45.3 ± 23.0% from a baseline points (postoperative OFF at 12 months) of 38.6 ± 12.9 to 21.1 ± 8.8 (DBS ON, Med OFF at same day). Disease duration at time of surgery was 10.4 ± 3.9 years and LEDD reduction was 52.8% (1072.72 in baseline vs. 484.57 at 12 months post surgery).

Preoperative baseline of the sample had been 32 ± 11 UPRDS-III points in Med OFF vs. 12 ± 5 points in Med ON conditions (53.5 ± 17.2% percent improvement in levodopa response).

### Image Registration

Co-registration results of T2- and if available PD- and FGATIR sequences as well as postoperative MR / CT to the anchor-modality (T1) were done using default presets (figure 2) and were accurate upon visual inspection. Results of the 3T example patient are shown in figure 3A. Similarly, fractional anisotropy (FA) computed from the preoperative dMRI acquisition in the 3T example patient was registered to T1. All preoperative sequences (including FA if available) were used for nonlinear registrations to template space in the ANTs-based approaches (shown in rows 1-3 in figure 4 for the 3T example). In these multispectral warps, the T1-scan was mapped to the T1 template, T2 to T2, PD to PD (figure 3B). The FA volume was instead paired with the FMRIB58 FA template (https://fsl.fmrib.ox.ac.uk/fsl/fslwiki/FMRIB58_FA) that had been registered to 2009b space using an MNI-152 6th-gen to 2009b space transform (Horn, 2016). Since no FGATIR-template exists in 2009b space, Lead-DBS automatically paired this scan with an aggregated PCA template (Horn, 2017). The linear three-step registration was included mainly for reproducibility purposes (Schönecker et al., 2009) but equally supports multispectral registrations. In the MAGeT Brain-like approaches (rows 4-5 in figure 4), only T1-, T2- and PD-weighted acquisitions were used given these sequences were available in the IXI database of age-matched peer-brains (i.e. “templates” in MAGeT nomenclature). SPM-based approaches (rows 6-7 in figure 4) used all preoperative acquisitions except the FA volume. Here, volumes were not paired with a specific template as in the ANTs-based registrations. Instead, tissue priors were used to learn posterior segmentations using voxel intensities across image modalities (Ashburner and Friston, 2005). All methods except the FNIRT and Linear Three-Step registrations were able to precisely segment the STN target region based on manual inspection. Note that the FNIRT method does not support multispectral warps and estimated the warp based on the T1 volume only (on which the STN is not visible). This may explain the mismatch in template vs. subject STN target regions.

**Figure 3:**
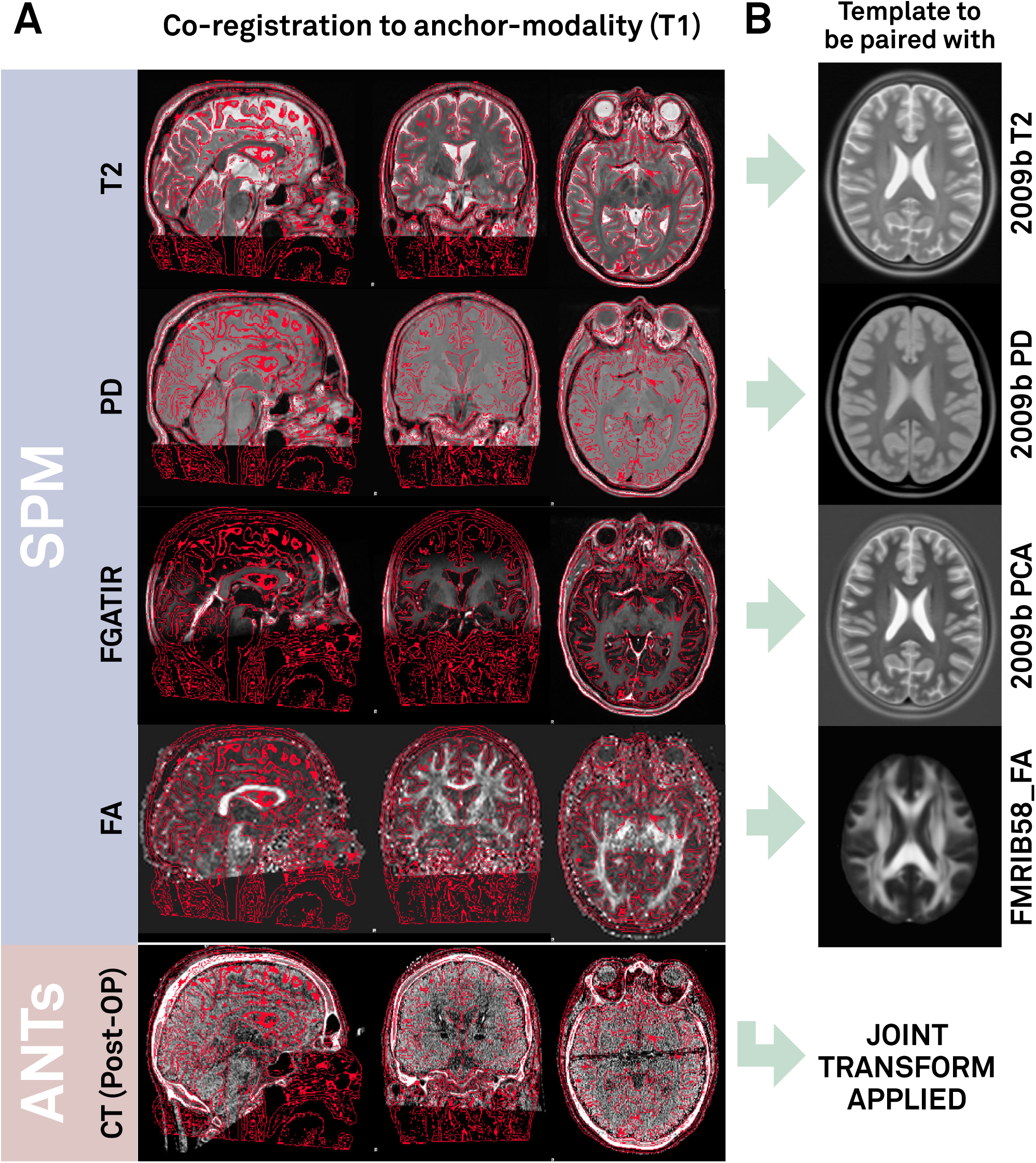
A) Co-registration results of the 3T example patient. Lead-DBS linearly registers preoperative T2, PD, FGATIR and FA volumes to the T1 anchor modality (visualized as red edge contours) using SPM. Similarly, by default, postoperative CT is linearly mapped to T1 using ANTs. A tone mapped version of the CT is shown (equally displaying brain- and bone-windows). B) In the following ANTs-based normalization step, the T1 volume will be registered to the T1-weighted MNI template (2009b NLIN Asym space; not shown). Likewise, T2 and PD volumes will be mapped to T2-/PD-templates. FGATIR volume by default is mapped to a synthetic PCA template while FA to a registered version of the FMRIB58_FA template. These five transforms result in a joint nonlinear deformation field that is equally applied to pre- and postoperative acquisitions.

**Figure 4:**
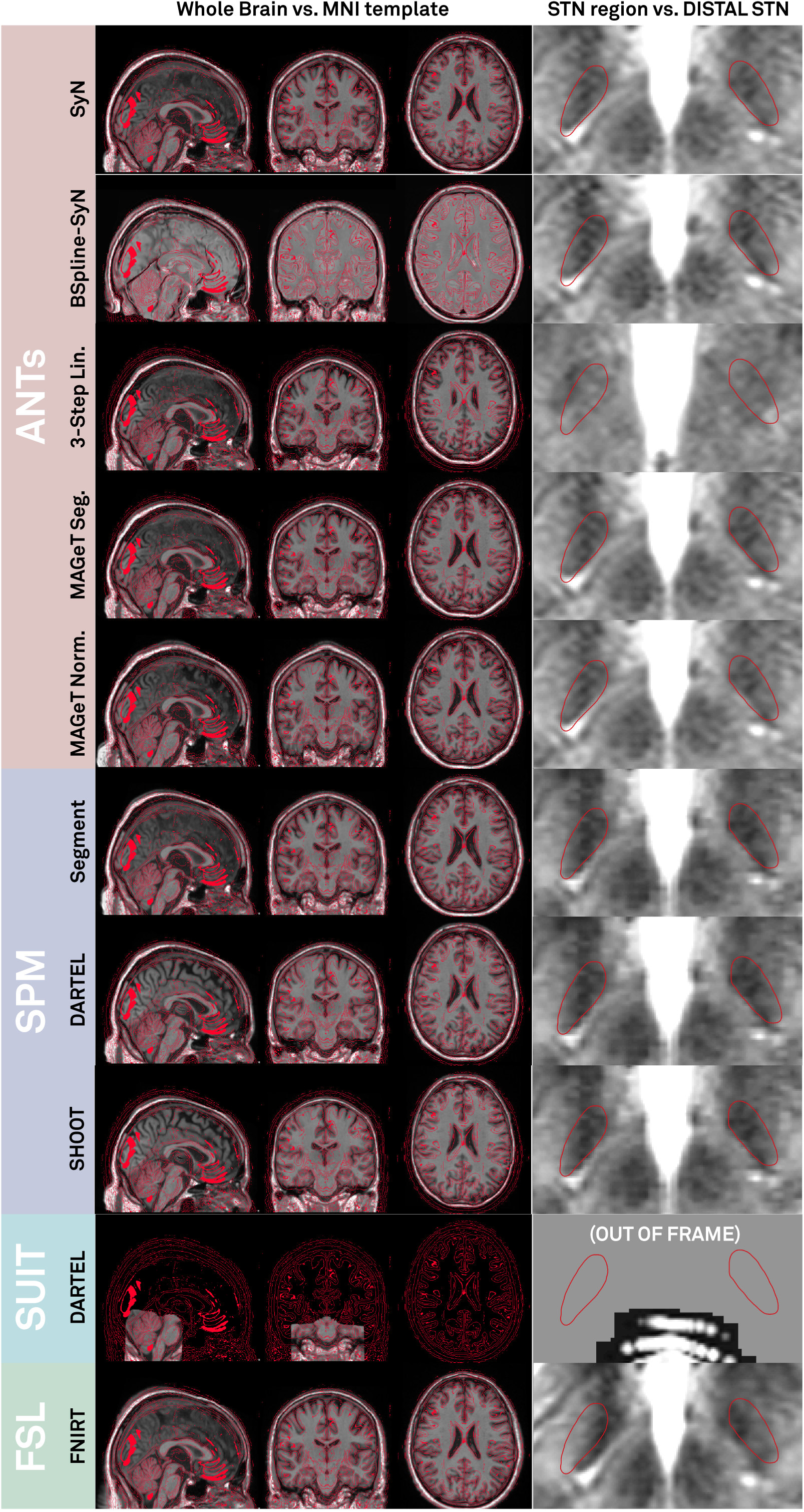
Normalization results of the 3T example patient. Based on the preoperative multimodal MRI (T1, T2, PD, FGATIR) of the patient, individual anatomy was registered into ICBM 2009b NLIN Asym (“MNI”) space using various methods. Left column: MNI space (red wireframes) overlaid to normalized T1 acquisition. Right column: DISTAL atlas STN (red wireframes) overlaid to normalized T2 acquisition. Note that the SUIT registration uses SPM methods too, but is based on a toolbox focusing on brainstem and cerebellum anatomy. Thus, normalizing the STN region with this preset is not possible, the method is still displayed for the sake of completeness. It is applicable for brainstem targets such as the Pedunculopontine Nucleus (PPN).

Finally, in the 3T example patient, brain shift correction led to a refined registration between postoperative CT and preoperative anchor modality (T1; figure 5). A shift of 0.17 mm to the left, 0.9 mm to anterior and 1.66 mm in dorsal direction was introduced (figure 4). In the present case, not much pneumocephalus was present and the example may rather demonstrate the introduction of higher robustness and precision with an additional subcortical refinement transform. An example of the tool in case of prevalent pneumocephalus may be seen in the methods figure S1.

**Figure 5:**
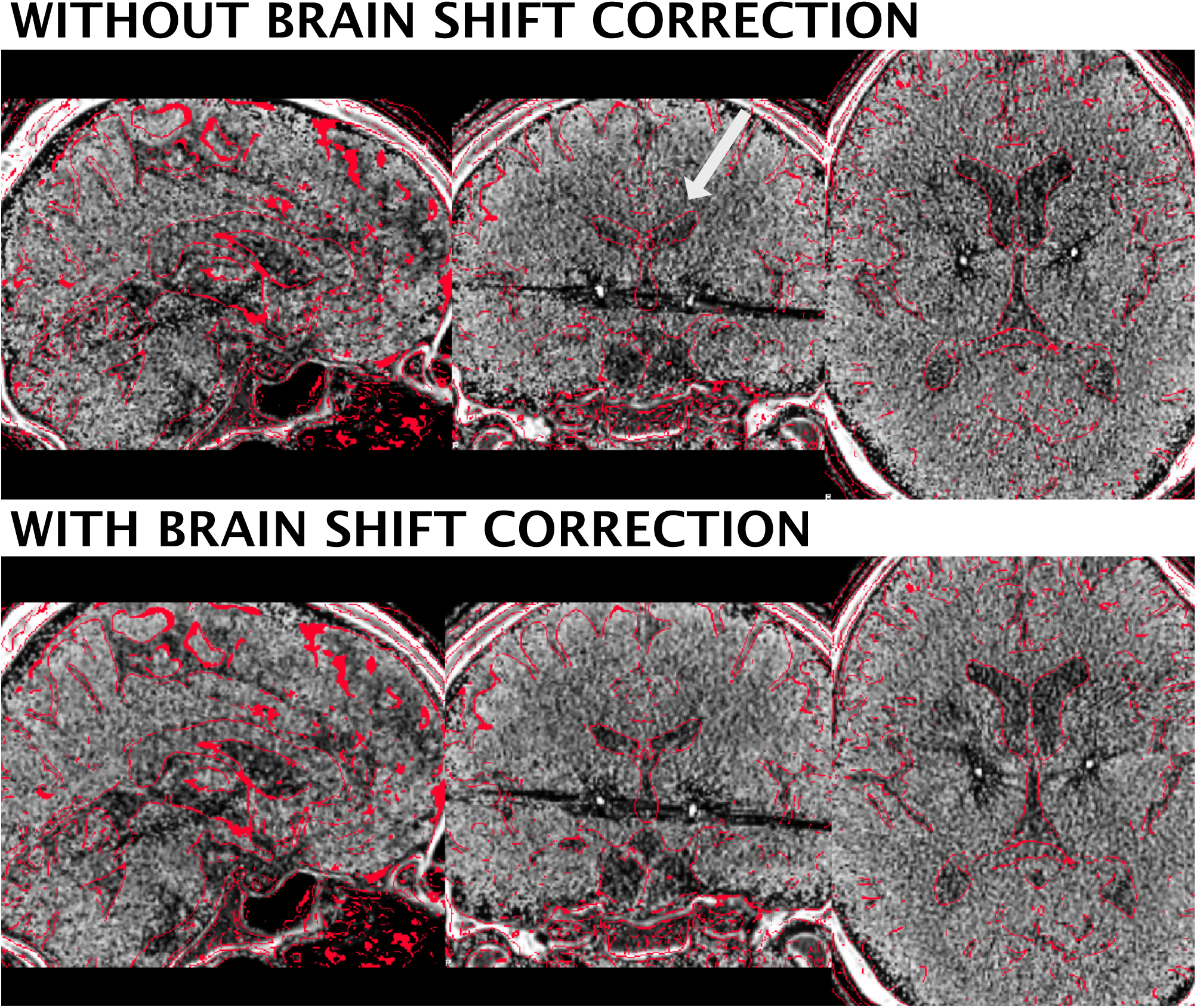
Brain shift correction results of the 3T example patient. The approach serves as a refinement registration step between post- and preoperative acquisitions and is able to minimize nonlinear registration errors due to pneumocephalus (figure S1). In the present example, the postoperative CT was shifted by 1.66 mm in z-, 0.9 mm in y- and 0.17 mm in x-direction. A better registration can best be seen in the area of the ventricles (white arrow). Postoperative CT was tone mapped to show contrast in both brain and skull windows.

### Electrode Reconstructions

In the 3T example patient, the PaCER method found an optimal solution including location of electrode contacts in a fully automated manner whereas the TRAC/CORE method robustly reconstructed the trajectory but contacts had to be adjusted manually. Final fully automatic PaCER reconstruction is shown in figure 6. In the subsequent step, orientation of the segmented electrode was reconstructed using the Directional Orientation Detection (DiODe) algorithm, an updated version of the approach described in (Sitz et al., 2017). Relative to a marker position pointing strictly to anterior, rotation of the electrodes was corrected by 65° (right lead) and 30° (left lead) clockwise as seen from the tip, respectively. Final MNI coordinates of planning (lowermost) contacts were x = 11.6, y = -16.2, z = -9.1 on the right and x = -12.7, y = -15.1, z = -10.7 mm on the left. Relative to the midcommissural point, in stereotactic coordinates, these corresponded to x = 11.5, y = -4.4, z = -5.9 (right) and x = -12.7, y = -3.9, z = -7.0 mm (left).

**Figure 6:**
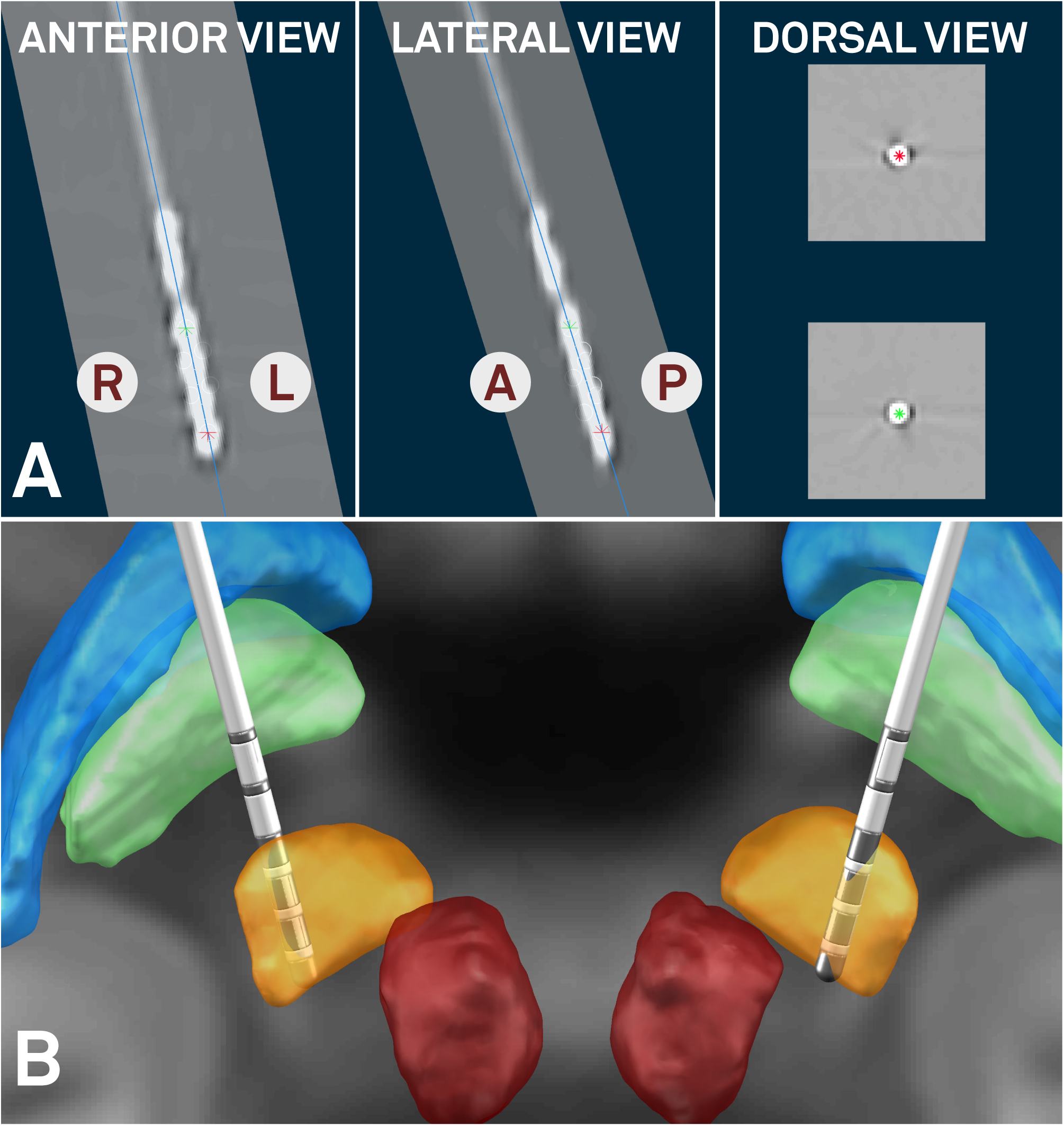
Fully automated electrode reconstruction results (PaCER method contributed by Husch and colleagues) of the 3T example patient. Orientation of lead reconstructed using the method by Sitz and colleagues. A) Postoperative CT is shown orthogonal to reconstructed trajectory (right hemisphere, blue line) in anterior, lateral and dorsal views. Ventral- and dorsalmost contacts marked by red and green asterisks, respectively. Using this view, users can fine-tune electrode reconstructions in a very precise way. B) Final 3D rendering of results in synopsis with key structures defined by the DISTAL atlas. Both electrodes placed in dorsolateral STN which corresponds to the sensorimotor functional zone of the STN. Right lead resides minimally more medial than left (in respect to atlas STN) which can be accounted for by field steering (figure 8).

### Microelectrode Recordings

After electrode reconstruction, classifications (no cell activity, unspecific cell activity, clear STN typic activity pattern) recorded in the 3T patient were mapped to the subcortex and are shown in figure 7 for the right hemisphere. On both sides, boundaries of firing patterns on the top and bottom of the STN corresponded well to the atlas-/imaging-defined STN. For instance, in the left lateral trajectory, no clear STN activity was reported and in agreement to that, the trajectory traverses outside (lateral) to the imaging-defined STN throughout its whole course.

**Figure 7:**
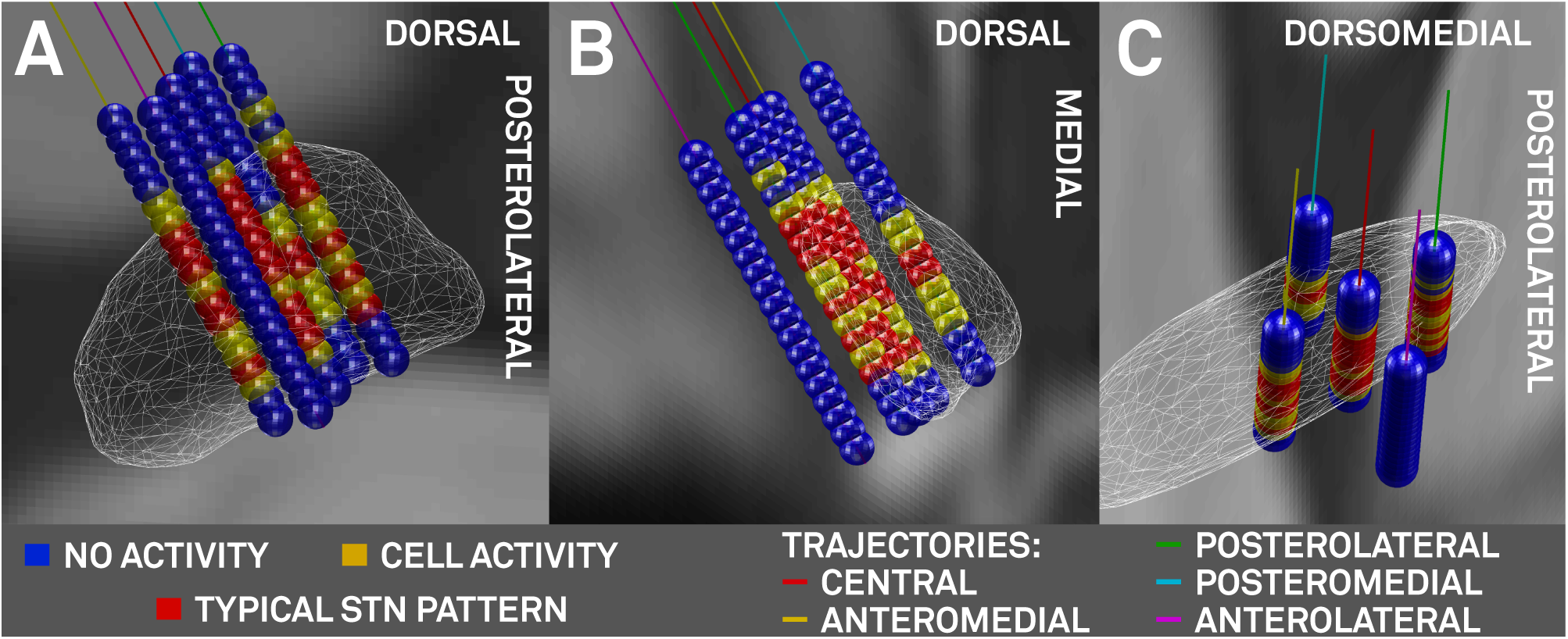
Left-Hemispheric microelectrode recording results of the 3T example patient. A) oblique view orthogonal to the lateral surface of the STN (white wireframes). B) view from posterior and C) dorsal. Markers in blue (no cell activity), yellow (cell activity), red (typical STN firing pattern) placed on 45° rotated Ben Gun (X-) array of microelectrodes between 7.5 and -1.5 mm distance to surgical target in 0.5 mm steps. Trajectories: central (red), lateral (magenta), medial (cyan), posterior (green) and anterior (yellow) trajectories.

### VTA Calculation

Stimulation parameters of 2 mA on the planning contact (ventral segmented level in ring mode) were calculated using the FEM-model and standard vs. frequency adapted conductivity values (figure 8 A, B). Here, the “heuristic” electric-field (E-field) threshold of 0.2 V/mm was used. The E-Field is the first derivative of the voltage distribution across the tissue surrounding the electrode. To demonstrate the possibilities of segmented electrodes, an additional unidirectional setting was calculated (figure 8 C). Finally, to allow for comparison with heuristic VTA models implemented in Lead-DBS, 2 V estimates on a ring electrode (Medtronic model 3389) are shown in figure 8 D, E using the Mädler or Kuncel models, respectively.

**Figure 8:**
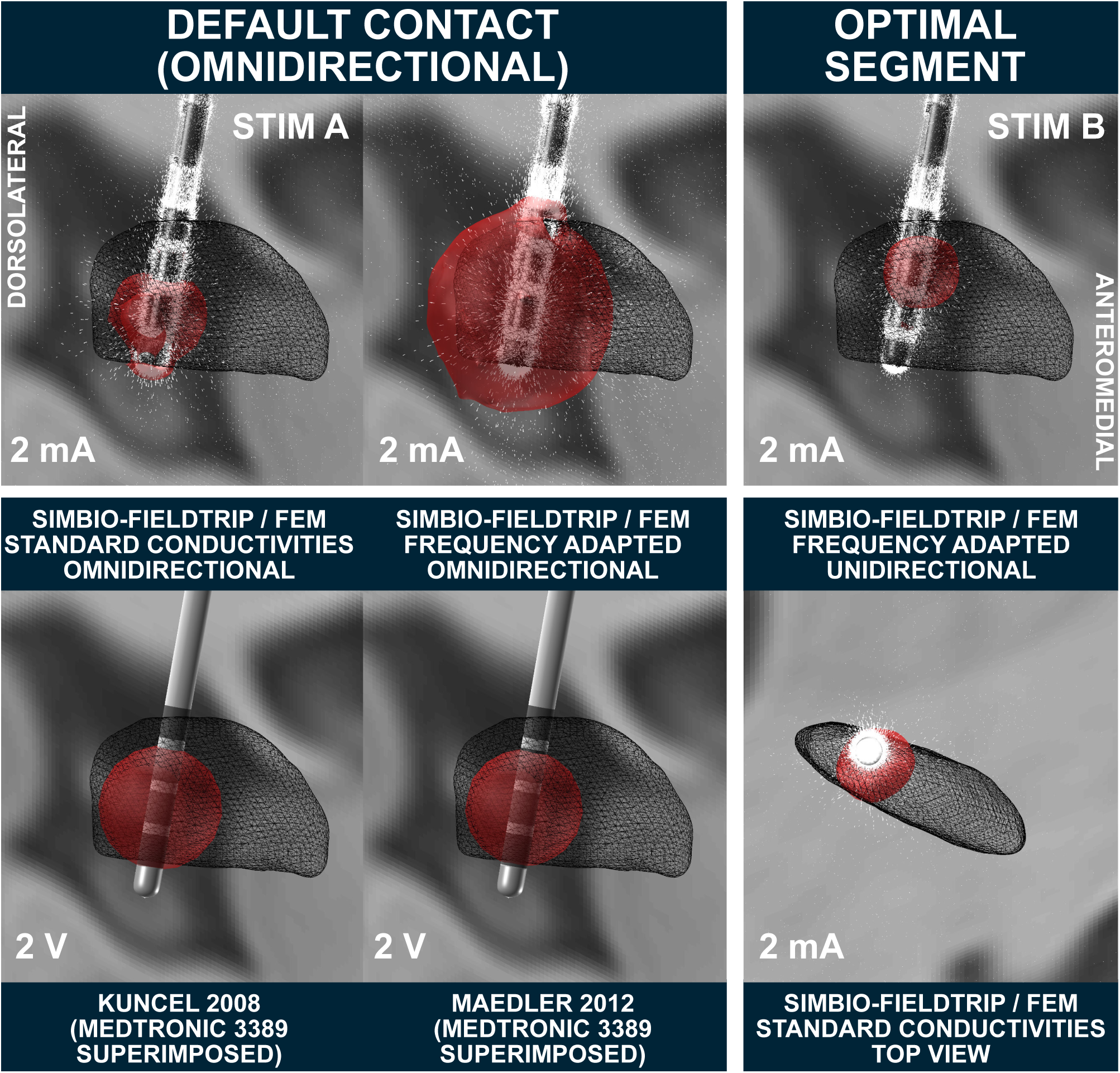
VTA modeling (right hemispheric lead) in the 3T example patient. Left two columns: several omnidirectional stimulations at the default contact. Top row demonstrates the strong impact on standard conductivities vs. frequency-adapted conductivities in the resulting FEM-based VTA. Bottom row shows two heuristic voltage driven models implemented in Lead-DBS. These models are not validated for directional leads, thus, a Medtronic 3389 electrode is visualized instead. Right column: The “optimal” segment on the top (K13) is used as cathode, steering the field anterolaterally to reach a good coverage of the dorsolateral STN. Simulations marked with Stim A and B are used as connectivity seeds in subsequent figures.

### Connectivity from VTA to other Brain Regions

Figure 9 A-C show results of the (3T patient‘s) patient-specific Generalized Q-sampling Imaging (GQI) and Gibbs‘-tracking approaches (whole connectome) as well as a Human Connectome Project (HCP) based normative connectome (Horn et al., 2017a; Setsompop et al., 2013). In figure 9 D-F, GQI based fiber tracts running through the VTA defined in figure 8A and C & D are shown. Figure 10 shows connectivity profiles from the VTA defined in figure 8A projected to the cortex using various structural and functional connectomes.

**Figure 9:**
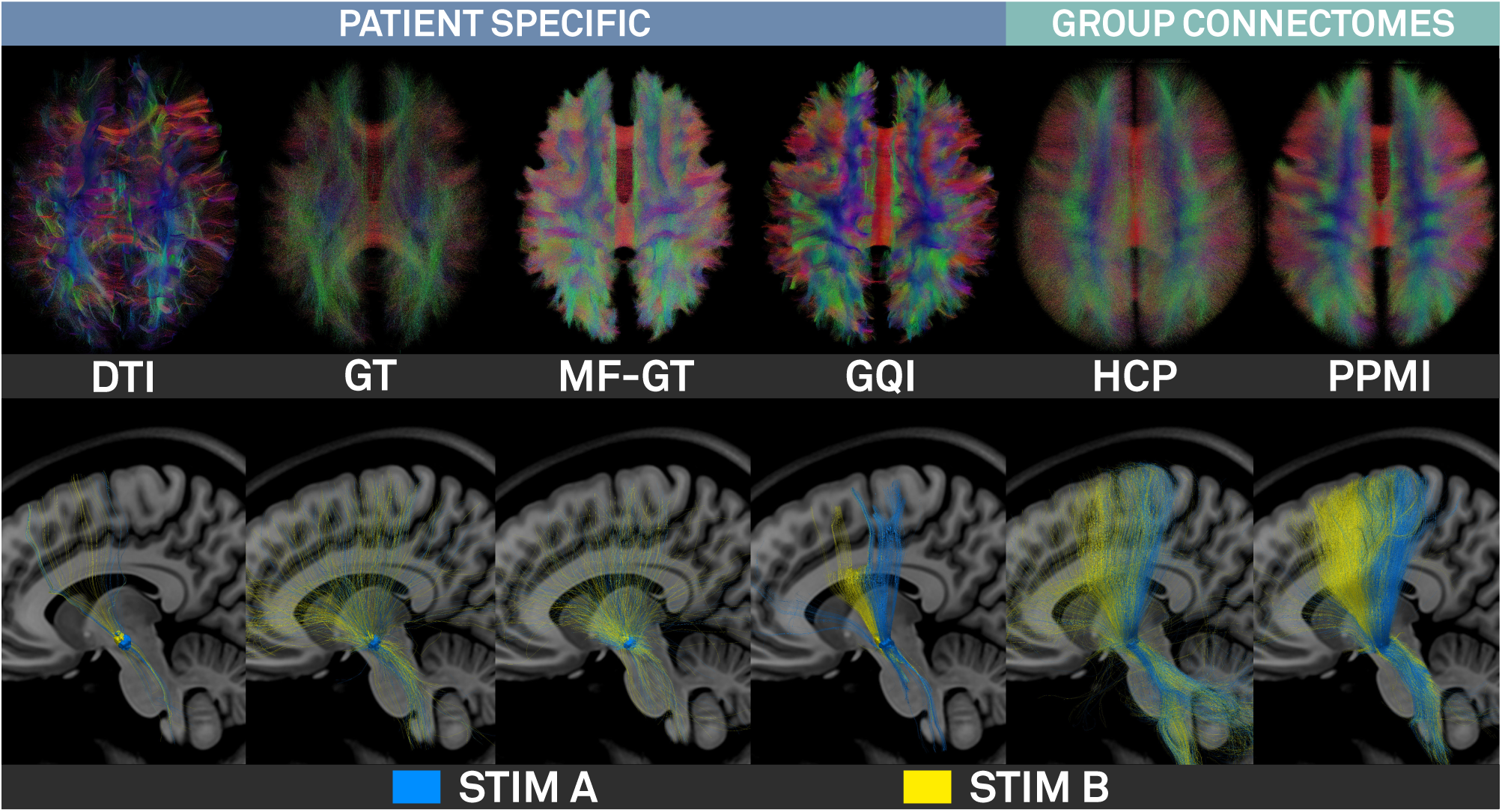
Fiber tracking results of the 3T example patient. First four columns show results of the patient-specific DTI, Gibbs’-tracking, Modelfree Meso global tracker and GQI approaches (top: whole connectome, bottom, tracts seeding from STIM A & B; see figure 8). Last two columns show the same views on HCP and PPMI based normative connectomes.

**Figure 10:**
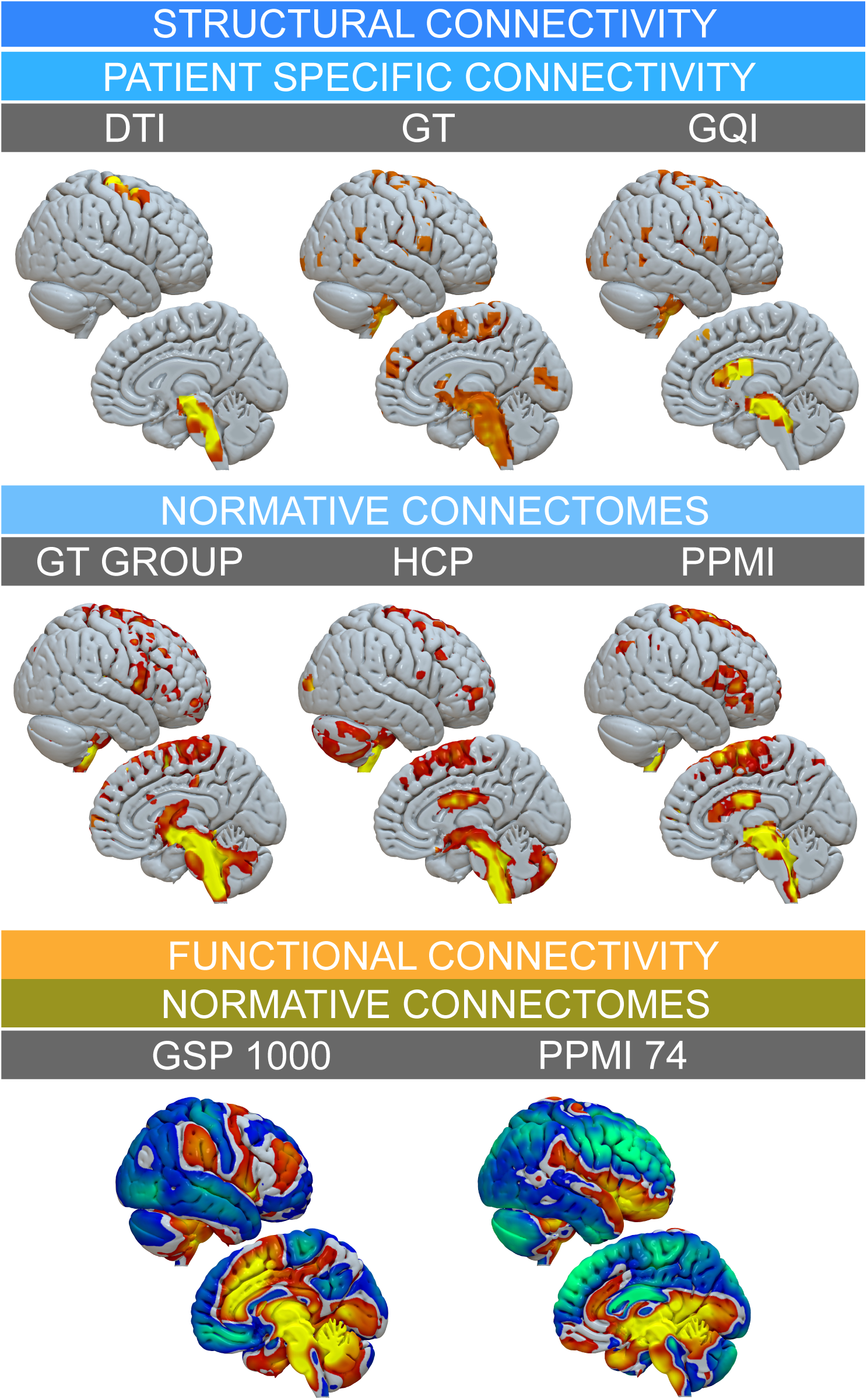
Connectivity from VTA defined by “STIM A” (figure 8) projected on the right hemispheric surface as defined by various connectomes. Top row: Patient specific structural connectivity using DTI, Gibbs’ Tracking and GQI methods. Mid row: Connections defined by the structural Gibbsconnectome, HCP Adult Diffusion and PPMI PD connectomes. Bottom row: Functional connectivity between VTA and other brain regions as defined by normative GSP 1000 and PPMI 74 PD connectomes. This figure demonstrates a multitude of options to analyze VTA connectivity in Lead-DBS, but also highlights challenges of the process, since different methods/datasets yield different results.

### Explaining clinical improvement in retrospective cohort

Results are summarized in table 9. Briefly, volumes of overlap correlated significantly with the empirical clinical outcome (FSL FNIRT: R = 0.38 at p = 0.007, ANTs SyN without / with subcortical refinement: R = 0.47 / 0.49 at p < 0.001; SPM New Segment: R = 0.52 at p < 0.001). Weighted overlaps between E-Field and STN correlated higher with clinical improvement for all normalization methods (FSL FNIRT: R = 0.46 at p < 0.001, ANTs SyN without / with subcortical refinement: R = 0.47 / 0.52 at p < 0.001 / < 10^-4^, SPM New Segment: R = 0.54 at p < 10^-4^). Mid column of figure 11 summarizes these findings. When adding additional clinical co-variates to a GLM to explain motor improvement (right column in figure 11), RMSE was comparable between methods (∼14-16%) but explained variance was 8% higher between best (SPM New Segment with E-Field) vs. worst (FSL FNIRT with binary VTA) methods (table 9).

**Figure 11:**
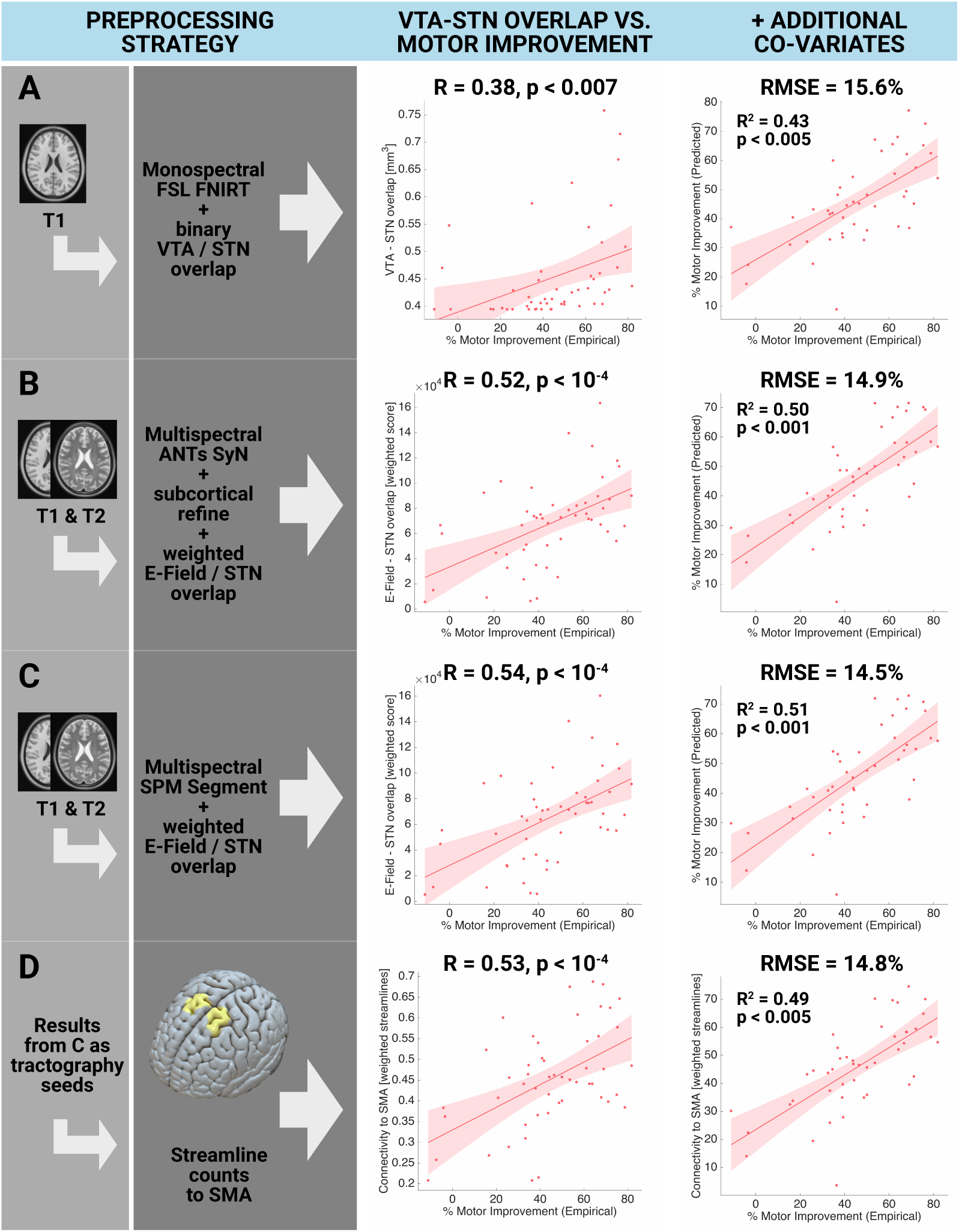
Amount of variance in clinical outcome explained when applying various preprocessing options (retrospective 1.5T cohort). A) an exemplary “standard neuroimaging” approach with a monospectral (T1 only) FSL FNIRT based registration and a binary VTA. B & C) Default pathways of Lead-DBS, registering preoperative data into template space in multispectral fashion according to the most optimal method as delineated in (Ewert et al. 2018a). In these approaches, the overlap sum between the E-Field gradient magnitude inside the STN was calculated. D) Using results from C as weighted seeds to isolate fibers from the normative PPMI connectome (table 8), correlating weighted numbers of streamlines to SMA to explain clinical motor improvement. First column of scatterplots shows direct correlation between VTA-STN overlap (weighted E-field overlap or weighted streamlines to SMA) with clinical improvement. Second column shows GLM with additional clinical covariates. Table 9 shows results for additional preprocessing strategies.

Panel A of figure 11 shows the FNIRT method (which does not work multispectrally and had only T1 MRIs as input, thus has poor if any information on the STN location) in combination with a binarized VTA. This approach could be seen as a “common neuroimaging” approach since most fMRI studies use T1 weighted images in their normalization step and binarized VTAs are common in the field. Panels B and C show the two multispectral Lead-DBS default pathways identified in (Ewert et al. 2018a) and apply a weighted VTA (E-Field magnitude). Both yield a similar outcome of ∼R = 0.5, increasing the amount of variance explained by imaging alone from ∼14% (FNIRT T1) to ∼26% (Lead-DBS defaults). Their connectivity strength (number of weighted streamlines to the SMA as defined by the 6ma entry – the medial and anterior parcel of sensorimotor numbered results – of the Glasser et al. 2016 parcellation) was calculated using Lead Connectome Mapper. Resulting values were equally correlated with empirical motor improvement scores (fig. 11 panel D).

## Discussion

We present a comprehensive and advanced processing pipeline to reconstruct, visualize and analyze DBS electrode placement based on neuroimaging data. Specific strengths in comparison to other tools are a seamless integration with a wide array of neuroimaging tools (table 11), a strong focus on precise spatial normalization and connection to a structural and functional connectome pipeline that facilitates connectivity analyses within the DBS context (figs. 9 & 10).

Contributions of the present paper are three-fold. First, an overview is provided regarding the novel neuroimaging methods that were added or updated over the course of four years since the initial publication of Lead-DBS. Second, a default pathway navigating through the multiple options in both DBS and connectome pipelines is outlined (figure 2). This pathway is motivated by both empirical data (Ewert et al., 2018a; Fillard et al., 2011; Klein et al., 2009; Maier-Hein et al., 2017) and by the experience of the Berlin DBS center where Lead-DBS or similar applications were used to localize roughly three thousand DBS electrodes since 2008. Third, results of various processing steps are visualized for a single patient and quantitatively analyzed in a retrospective analysis of 51 patients. In the latter, we do not only demonstrate that overlap between stimulation volumes and the subthalamic nucleus may explain clinical motor improvement, but we also show that the amount of variance explained may depend on the applied preprocessing strategy. Specifically, in the cohort investigated here, the advanced multispectral normalization pipelines implemented as defaults in Lead-DBS are able to explain more variance in clinical outcome than a “typical neuroimaging” pipeline.

In total, Lead-DBS includes five methods for linear co-registrations (table 1), ten normalization approaches (table 3), four approaches for electrode reconstructions, four VTA models (table 4) and four whole-brain tractography pipelines (table 6). Twenty-four subcortical atlases (table 2) and 17 brain parcellations (table 7) are pre-installed. Finally, two functional and four structural connectomes were converted into a format suitable for use in Lead-DBS (table 8). Taken together, these resources build a comprehensive toolbox for DBS electrode localization and the analysis of local (coordinate- or VTA-based) and global (structural and functional connectivity) features. While the number of these methods may introduce complexity, a user-friendly “default pathway” (figure 2) was established which works robustly and well for most applications. This pathway was established while working on several studies that were empowered by Lead-DBS with a variety of clinical and scientific aims. Some of these used Lead-DBS to integrate population based neural activity with anatomical structures (“Subcortical Electrophysiology Mapping” approach; Accolla et al., 2017; Geng et al., 2018; Horn et al., 2017b; Lofredi et al., 2018; Neumann et al., 2017; van Wijk et al., 2017). Other studies used connectivity profiles from DBS electrodes to predict clinical outcome (Horn et al., 2017c) or combined electrophysiological measures with DBS contact connectivity profiles (Accolla et al., 2016). In an effort to improve the safety profile of DBS implantations, some aimed at determining the relationship between electrode positions and clinical side effects or non-motor symptoms (Mosley et al., 2018b; 2018a). Finally, in other publications, the main aim was to ensure that the analyzed electrodes were indeed placed within the target region (Barow et al., 2014; Brücke et al., 2014; Ehlen et al., 2017; Hohlefeld et al., 2015; 2017; 2013; Krause et al., 2015; 2016; Kroneberg et al., 2017; Merkl et al., 2016; Neumann et al., 2015; 2016; Schroll et al., 2015; Tiedt et al., 2016).

**Table 6:** Brain parcellations (of use for connectomic analyses) suitable for / available within Lead-DBS.

| Atlas Name | Pre-installed in Lead-DBS | Source(s) of Information | Structures of Focus / Specialities | Publication(s) |
| --- | --- | --- | --- | --- |
| **Whole Brain** |  |  |  |  |
| Harvard-Oxford cortical/subcortical atlases | + | Manual segmentations | Whole Brain | Makris et al. 2006, Frazier et al. 2005, Desikan et al. 2006, Goldstein et al. 2007 |
| MICCAI 2012 Multi-Atlas Labeling Workshop and Challenge (Neuromorphometrics) | + | Manual segmentations | Whole Brain | neuromorphometrics.com |
| Brainnetome Atlas parcellation | + | fMRI, dMRI | Whole Brain | Fan et al. 2016 |
| Automated Anatomical Labeling (v2) | + | MRI | Whole Brain | Tzourio-Mazoyer et al. 2002 |
| AICHA: An atlas of intrinsic connectivity of homotopic areas | + | fMRI | Whole Brain | Joliot et al. 2015 |
| Hammers_mith Atlas | + | MRI | Whole Brain, special focus on temporal lobe | Hammers et al. 2003, Gousias et al. 2008 |
| PrAGMATIC | – | fMRI | Functional atlas based on task fMRI | Huth et al. 2016, Huth et al. 2015 |
| fMRI-based random parcellations | + | fMRI | Fine-grained random parcellations informed by rs-fMRI data | Craddock et al. 2012 |
| Voxelwise parcellations | + | MRI | Whole Brain | Horn & Kühn 2015 |
| **Cortex Focus** |  |  |  |  |
| Mindboggle 101 | + | Manual segmentations | Cortex / Desikan protocol | Klein et al. 2012 |
| Yeo functional parcellations | + | fMRI | Cortex | Yeo et al. 2011 |
| Local-Global Parcellation of the Human Cerebral Cortex | + | fMRI | Cortex, refining Yeo 2011 parcellations | Schaefer et al. 2017 |
| Cortical Area Parcellation from Resting-State Correlations | + | rs-fMRI | Cortex | Gordon et al. 2016 |
| HCP MMP 1.0 | – | Multimodal | Cortex, surface maps | Glasser et al. 2016 |
| Desikan-Killiany Atlas | – | Gyrification related ROI | Cortex, surface maps | Desikan et al. 2006 |
| Destrieux Atlas | – | Gyrification related ROI | Cortex, surface maps | Destrieux et al. 2010, Fischl et al. 2004 |
| MarsAtlas | – | Gyrification related ROI | Cortex, surface maps | Auzias et al. 2016, Auzias et al. 2013 |
| **Specific Subregions** |  |  |  |  |
| JuBrain / Juelich histological atlas | + | Histology | Specific regions | Zilles et al. 2010, Amunts et al. 2007, Eickhoff et al. 2005, Eickhoff et al. 2010, Eickhoff et al. 2006bp |
| Human Motor Area Template | + | fMRI / Metaanalysis | (Pre-) motor cortex | Mayka et al. 2006 |
| Sensorimotor Area Tract Template | + | dMRI | Corticospinal tract subregions | Archer et al. 2017 |
| SUIT cerebellar parcellation | + | MRI | Cerebellum | Diedrichsen et al. 2006, Diedrichsen et al. 2011 |
| Buckner functional cerebellar parcellation | + | fMRI | Same networks as in Yeo 2011 cortical parcellations defined in cerebellum | Buckner et al. 2011 |

**Table 7:** Electrode models included in Lead-DBS.

| Manufacturer | Type | # Contacts | Contact Spacing | Omnidirectional/<br>Segmented |
| --- | --- | --- | --- | --- |
| DBS |  |  |  |  |
| Medtronic | 3389 | 4 | 2 mm | O |
|  | 3387 | 4 | 3 mm | O |
|  | 3391 | 4 | 7 mm | O |
| Boston Scientific | Vercise | 8 | 2 mm | O |
|  | Vercise Directed | 8 | 2 mm / Segmented | S |
| Abbott / St. Jude Medical | Active Tip 6146-6149 | 4 | 2 mm | O |
|  | Active Tip 6142-6145 | 4 | 3 mm | O |
|  | Infinity Directed 6172 | 8 | 2 mm / Segmented | S |
|  | Infinity Directed 6173 | 8 | 2 mm / Segmented | S |
| PINS Medical | L301 | 4 | 2 mm | O |
|  | L302 | 4 | 3 mm | O |
|  | L303 | 4 | 6 mm | O |
| iEEG |  |  |  |  |
| SDE | 08 S8 | 8 | 3.5 mm | O |
|  | 08 S10 | 10 | 3.5 mm | O |
|  | 08 S12 | 12 | 3.5 mm | O |
|  | 08 S16 | 16 | 3.5 mm | O |

**Table 8:** Normative Connectomes available in Lead-DBS format.

| Name | Repository | N | Population | Publication – Dataset | Processing Methods | Publication – Processing |
| --- | --- | --- | --- | --- | --- | --- |
| rs-fMRI based “functional cor |  |  |  |  |  |  |
| Functional group connectome 1000 healthy subjects GSP | Harvard Genomic Superstruct Project (GSP) | 1000 | Controls | Yeo et al. 2011 | Lead Connectome fMRI pipeline | Horn et al. 2017 |
| Functional group connectome 74 PPMI PD-patients, 15 controls | Parkinson’s Disease Progression Marker Initiative (PPMI) | 74 / 15 | PD patients, controls | Marek et al. 2011 | Lead Connectome fMRI pipeline | Horn et al. 2017 |
| dMRI based “structural conne |  |  |  |  |  |  |
| Structural group connectome 20 subjects Gibbs-tracker | Horn 2014 study | 20 | Controls | Horn et al. 2014 | Lead Connectome, Gibbs’ tracker | Horn et al. 2014 |
| Structural group connectome 169 NKI subjects Gibbs-tracker | Horn 2016 study | 169 | Controls | Horn et al. 2016 | Lead Connectome, Gibbs’ tracker | Horn et al. 2016 |
| Structural group connectome 32 Adult Diffusion HCP subjects GQI | HCP MGH Adult Diffusion dataset | 30 | Controls | Setsompop et al. 2013 | Lead Connectome, DSI Studio (GQI) | Horn et al. 2017 |
| Structural group connectome 90 PPMNI PD-patients GQI | Parkinson’s Disease Progression Marker Initiative (PPMI) | 90 | PD patients | Marek et al. 2011 | Lead Connectome, DSI Studio (GQI) | Ewert et al. 2018 |

**Table 9:** Preprocessing strategies used to explain variance in 1.5T retrospective cohort.

| VTA strategy |  | MR volumes used | Shown in fig. 11 | Correlation (image metric vs. motor improvement) |  | GLM (+ additional covariates) |  |  |  |
| --- | --- | --- | --- | --- | --- | --- | --- | --- | --- |
| Normalization strategy |  |  |  | R | p | RSME [%] | R<sup>2</sup> | Overall p | Significant predictors (p < 0.05) |
| **Binary VTA overlap** |  |  |  |  |  |  |  |  |  |
| Monospectral FSL FNIRT |  | T1 | Panel A | 0.38 | < 0.007 | 15.1 | 0.43 | < 0.005 | Sex |
| Multispectral ANTs SyN | w/o subcortical refine | T1 & T2 | – | 0.46 | < 0.001 | 15.3 | 0.45 | < 0.005 | Sex |
|  | with subcortical refine | T1 & T2 | – | 0.49 | < 0.001 | 14.8 | 0.48 | < 0.005 | Sex, VTA overlap |
| Multispectral SPM New Segment |  | T1 & T2 | – | 0.51 | < 10<sup>-4</sup> | 14.7 | 0.49 | < 0.001 | Sex, VTA overlap |
| **Weighted E-field VTA / STN overlap** |  |  |  |  |  |  |  |  |  |
| Monospectral FSL FNIRT |  | T1 | – | 0.46 | < 0.001 | 15.1 | 0.46 | < 0.005 | Sex |
| Multispectral ANTs SyN | w/o subcortical refine | T1 & T2 | – | 0.47 | < 0.001 | 15.3 | 0.46 | < 0.005 | Sex |
|  | with subcortical refine | T1 & T2 | Panel B | 0.52 | < 10<sup>-4</sup> | 14.9 | 0.48 | < 0.005 | Sex, E-Field overlap |
| Multispectral SPM New Segment |  | T1 & T2 | Panel C | 0.54 | < 10<sup>-4</sup> | 14.5 | 0.51 | < 0.001 | Sex, E-Field overlap |
| **Weighted streamline counts to SMA** |  |  |  |  |  |  |  |  |  |
| Weighted streamline counts to SMA seeding from E-Field (based method one row above) |  | T1 & T2 | Panel D | 0.53 | < 10<sup>-4</sup> | 14.8 | 0.49 | < 0.005 | Sex, Streamlines to SMA |

### The precision of Lead-DBS

Quantifying the precision of the processing pipeline is a difficult task but a frequently asked question and crucial to the widespread use of the tool. Unfortunately, without postmortem histological examination, no real ground truth exists. However, some indirect measures may help to address the question. First, it should be mentioned that errors may originate from several sources including i) MR-distortion artifacts, ii) within-patient co-registration including brain shift, iii) patient-to-template normalization and iv) electrode localization. Quantifying the first source falls under the domain of MR physics research and goes beyond the present scope. Still, it is advisable to apply distortion correction steps in each MR sequence if possible – even more so when high field magnets are involved. Second, errors in linear co-registration can be minimized by care- and skillful inspection of the data. The check co-registation and brain shift correction modules were specifically designed for the task at hand and iteratively improved to suit the needs and precision of DBS imaging. For instance, the brain shift correction step often sensibly corrects registrations on a submillimeter scale (figs 5 and S1). Normalization procedures were recently addressed in a comparative study (Ewert et al., 2018a). The study defined the default normalization pipeline which, depending on data quality, resulted in an average surface distance of STN/ GPi boundaries between 0.38 and 0.75 mm, while inter-rater distance was between 0.41 and 0.82 mm. Based on these results, the pipeline is able to segment STN and GPi nuclei equally well as human experts. In this context, an anteroposterior iron gradient in the STN poses an additional problem for this specific target. The posterolateral sensorimotor part of the nucleus that is targeted to treat movement disorders contains less iron than its anteriomedial parts, often rendering it smaller on MRI than it actually is (Dormont et al. 2004, Richter et al. 2004, Schäfer et al. 2011, de Hollander et al. 2014, Massey et. al 2012). This makes the registration of this specific nucleus to a template space more error-prone than other targets and yet again raises the need for ultra high-field multispectral preoperative imaging (e.g. see Forstmann et al. 2017, Keuken et al. 2014). Fourth, in the electrode reconstruction step, prior studies have used phantoms to verify that electrode artifact centers in MRI (Pollo et al., 2004; Yelnik et al., 2003) and CT (Hemm et al., 2009; Husch et al., 2017) indeed correspond to electrode centers in the brain. To this end, the PaCER algorithm as the default reconstruction algorithm for postoperative CT yielded an average reconstruction error below 0.2 mm – again depending on data quality. On the other end of the spectrum, the “x-ray mode” of Lead-DBS was specifically designed to reduce errors introduced by partial volume effects in imaging data of suboptimal resolution. In summary, all sources of error can be minimized by using high-quality imaging data, distortion correction and careful inspection or registration and localization results. Based on the retrospective cohort analysed in the present study, we demonstrate that the specialized and elaborate default pipeline of Lead-DBS may add to the amount of variance explained in DBS imaging data.

### Reproducibility, Open Science & Experimental Features

As stated above, a key mission of Lead-DBS development is to provide a platform for DBS imaging that is and remains i) free of use, ii) reproducible, open source & transparent, and iii) independent from commercial manufacturers. While this hinders the application in a clinical context (see below), within research, it has several advantages. First, the free software nature offers excellent worldwide accessibility, the possibility of fast skill dissemination in open workshops or courses within the academic field. Similarly, it is easy to script, automate and modify as permitted by the open source license while this is tedious or impossible in closed environments of clinical software. Second, transparent and open source code that is developed in a version controlled fashion (https://github.com/leaddbs/leaddbs) permits excellent reproducibility that is required for good scientific practice. In contrast, undocumented changes or discontinuity in the commercial applications may impose risks for producing consistent results. Discontinuation of commercial products has happened on multiple occasions in the field of DBS (table 10). Consequently, published studies that used discontinued software exist and are now hard if not impossible to reproduce. A slightly less obvious advantage of academic software is that its development is much more flexible. Commercial applications undergo highly involved and time-consuming certification processes to achieve CE-marks or FDA-approvals for safe use in clinical context. Needless to say, this is a great advantage or even requirement for clinicians but may drastically slow down software development. Furthermore, new research tools may not be easily integrated into commercial pipelines since these would require re-certification. In contrast, new tools can be integrated into academic software from idea and concept to end-user deployment within days. For instance, in 2009, the global fiber tracking approach (Reisert et al., 2011) won the Neurospin Fiber Cup evaluated as the best fiber tracking software compared to nine competitors (Fillard et al., 2011). With help from its developers, the Gibbs’ Tracker was integrated into the Lead Connectome pipeline within weeks. Recently, a newer comparative study found that the generalized q-sampling algorithm implemented in DSI studio yielded the highest “valid connection” score (Maier-Hein et al., 2017). Again, with kind support and permission of the developer, this method was integrated into Lead-DBS. A last example is the brain shift correction feature that was developed from idea to published code during a three day “brainhack global” event in 2017 at MIT (Craddock et al., 2016). Finally, a strong focus of clinical applications lies on their usability and processing speed. This is important since tools are used by medical personnel working under stressful circumstances where introduction into various complex software tools and long processing times are not tolerable. In contrast, in a research setting, search for innovative application, and development of new features outweighs the burden of computational time. Often, high performance compute clusters are available or jobs are run overnight. Thus, processes with high computational cost will be optimized for speed and standard applications in the former and for development and precision in the latter context. These thoughts illustrate that both types of tools – i.e. clinical vs. academic software – are needed. Given contradictory demands, a one-stop solution serving all purposes is hard if not impossible to create.

**Table 10:**
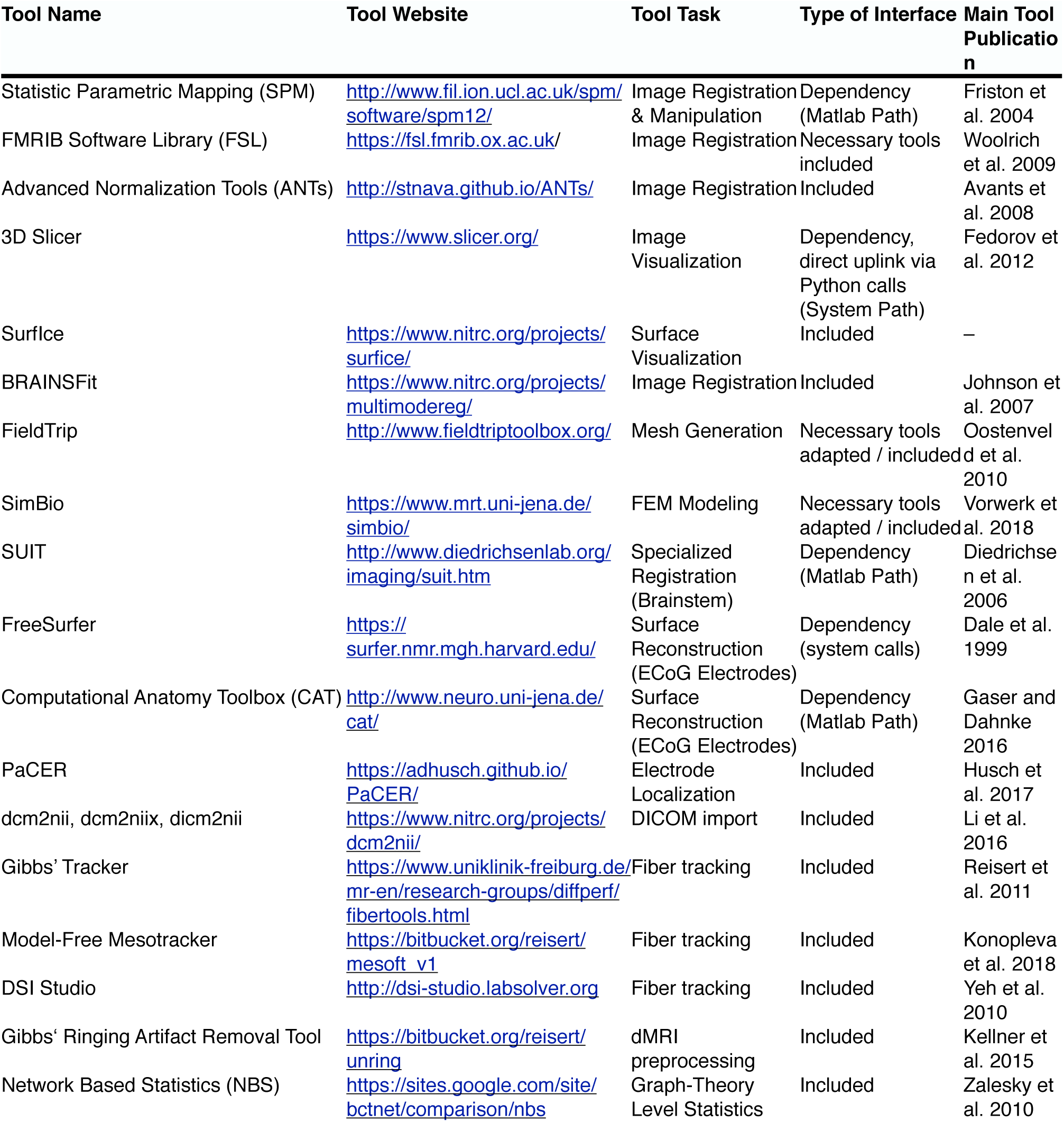
Interfaces between Lead-DBS and other neuroimaging tools. Except for FreeSurfer, all tools are available on Windows, macOS and Linux. Necessary binaries for FSL tools were adapted to work on Windows although there is no official Windows FSL release.

| Tool Name | Tool Website | Tool Task | Type of Interface | Main Tool Publication |
| --- | --- | --- | --- | --- |
| Statistic Parametric Mapping (SPM) | http://www.fil.ion.ucl.ac.uk/spm/software/spm12/ | Image Registration & Manipulation | Dependency (Matlab Path) | Friston et al. 2004 |
| FMRIB Software Library (FSL) | https://fsl.fmrib.ox.ac.uk/ | Image Registration | Necessary tools included | Woolrich et al. 2009 |
| Advanced Normalization Tools (ANTs) | http://stnava.github.io/ANTs/ | Image Registration | Included | Avants et al. 2008 |
| 3D Slicer | https://www.slicer.org/ | Image Visualization | Dependency, direct uplink via Python calls (System Path) | Fedorov et al. 2012 |
| SurfIce | https://www.nitrc.org/projects/surfice/ | Surface Visualization | Included | – |
| BRAINSFit | https://www.nitrc.org/projects/multimodereg/ | Image Registration | Included | Johnson et al. 2007 |
| FieldTrip | http://www.fieldtriptoolbox.org/ | Mesh Generation | Necessary tools adapted / included | Oostenveld et al. 2010 |
| SimBio | https://www.mrt.uni-jena.de/simbio/ | FEM Modeling | Necessary tools adapted / included | Vorwerk et al. 2018 |
| SUIT | http://www.diedrichsenlab.org/imaging/suit.htm | Specialized Registration (Brainstem) | Dependency (Matlab Path) | Diedrichsen et al. 2006 |
| FreeSurfer | https://surfer.nmr.mgh.harvard.edu/ | Surface Reconstruction (ECoG Electrodes) | Dependency (system calls) | Dale et al. 1999 |
| Computational Anatomy Toolbox (CAT) | http://www.neuro.uni-jena.de/cat/ | Surface Reconstruction (ECoG Electrodes) | Dependency (Matlab Path) | Gaser and Dahnke 2016 |
| PaCER | https://adhusch.github.io/PaCER/ | Electrode Localization | Included | Husch et al. 2017 |
| dcm2nii, dcm2niix, dicm2nii | https://www.nitrc.org/projects/dcm2nii/ | DICOM import | Included | Li et al. 2016 |
| Gibbs' Tracker | https://www.uniklinik-freiburg.de/mr-en/research-groups/diffperf/fibertools.html | Fiber tracking | Included | Reisert et al. 2011 |
| Model-Free Mesotracker | https://bitbucket.org/reisert/mesoft_v1 | Fiber tracking | Included | Konopleva et al. 2018 |
| DSI Studio | http://dsi-studio.labsolver.org | Fiber tracking | Included | Yeh et al. 2010 |
| Gibbs' Ringing Artifact Removal Tool | https://bitbucket.org/reisert/unring | dMRI preprocessing | Included | Kellner et al. 2015 |
| Network Based Statistics (NBS) | https://sites.google.com/site/bctnet/comparison/nbs | Graph-Theory Level Statistics | Included | Zalesky et al. 2010 |

**Table 11:**
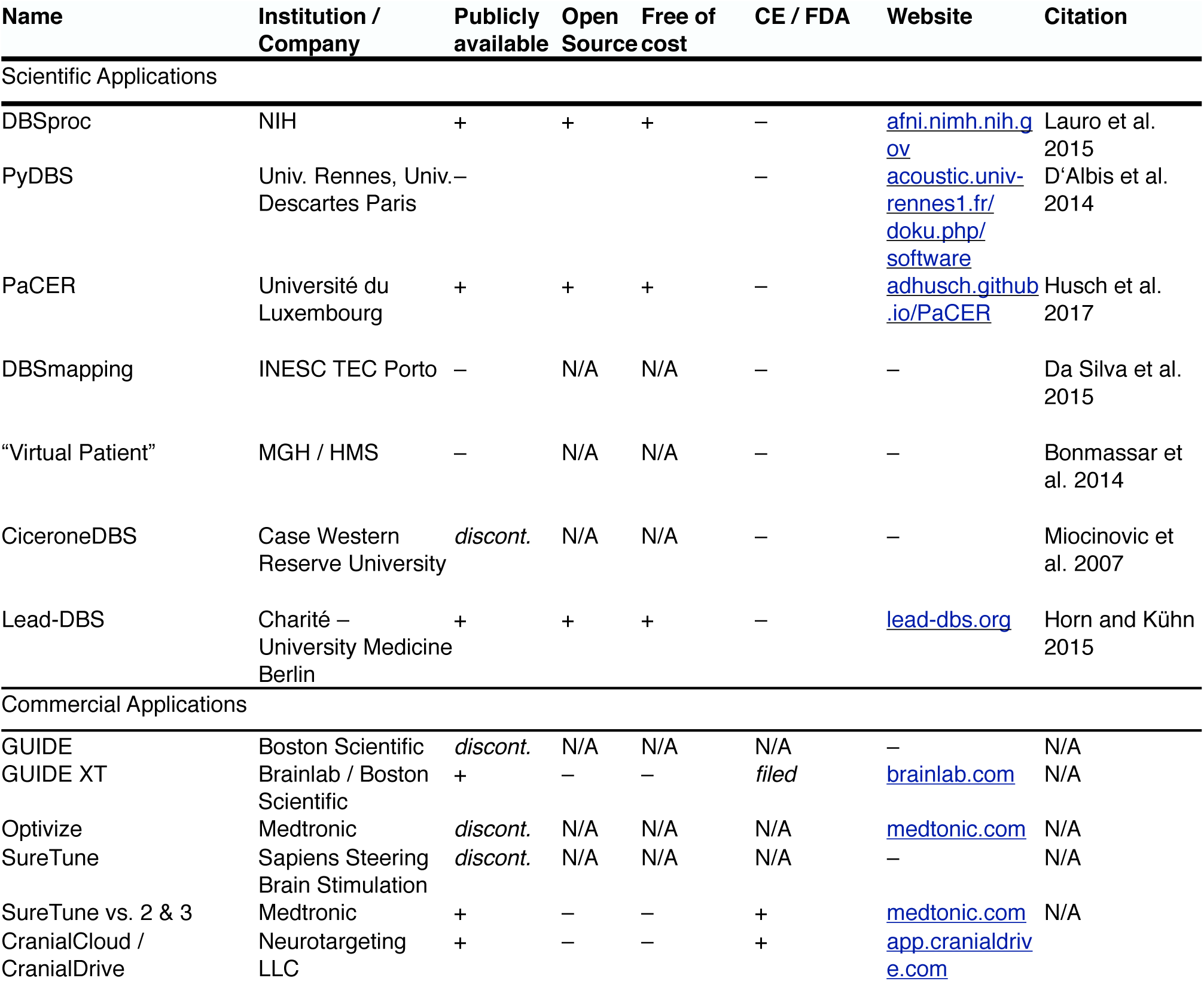
Software with aims comparable to Lead-DBS. Entries based on publicly available information at the time of writing.

## Limitations and Future Directions

In comparison to the first release, version 2 represents a major update and a drastically enhanced pipeline for DBS imaging. However, further development is planned to address remaining limitations and further maximize precision of the pipeline. To this end, the pipeline and resulting models may be broken down into four layers, each of which could be further improved as technology and methodology advance (figure 12). First, an anatomy layer describes the local surroundings of the electrode and helps to define electrode placement initially. This layer is presently defined by imaging and brain atlases (of which some may be informed by histology or other sources of information, table 2). It was mentioned multiple times that naturally, overall precision drastically depends on imaging quality (Ewert et al., 2018a; Husch et al., 2017). Crucially, the MR protocol of the 3T example patient represents a state-of-the art pipeline achievable in typical hospital settings and comprises a specialized basal ganglia sequence (FGATIR; Sudhyadhom et al., 2009). However, the diffusion-MRI acquired here may not be optimally suited to investigate the fine and complex details around DBS targets but was possible to scan within clinical routine. An example of a more optimal scan protocol can be found in (Akram et al., 2017). Moreover, as discussed in our original article (Horn & Kühn 2015), the use of postoperative CT or MRI each has specific advantages (higher signal to noise of the electrode on CT but direct visibility of surrounding anatomical structures on MRI, no radiation). It is hard to tell on empirical grounds which is better but the visibility of structures on the MRI generates a strong argument in favor for postoperative MRI – with the possibility to much more deliberately control for accuracy of post- to pre-co-registration around the target region. Similarly specialized methods like quantitative susceptibility mapping (Wang and Liu, 2015) or the use of ultra-highfield MRI (Forstmann et al., 2017; 2014) are other potential ways of increasing anatomical precision. As figure 1 illustrates, DBS target regions are typically small in size and reside in complex surroundings with a multitude of fiber tracts and functional segregations. Thus, sources above and beyond MRI may be needed to refine definition on the anatomy layer. To this end, techniques like polarized light imaging (Axer et al., 2011) or anisotropic scattering imaging (Shin et al., 2014) as well as the registration of histological stacks into template space (Alho et al., 2017; Amunts et al., 2013; Chakravarty et al., 2006; Ewert et al., 2018b; Forstmann et al., 2016; Jakab et al., 2012; Yelnik et al., 2007) are already applied increasingly.

**Figure 12:**
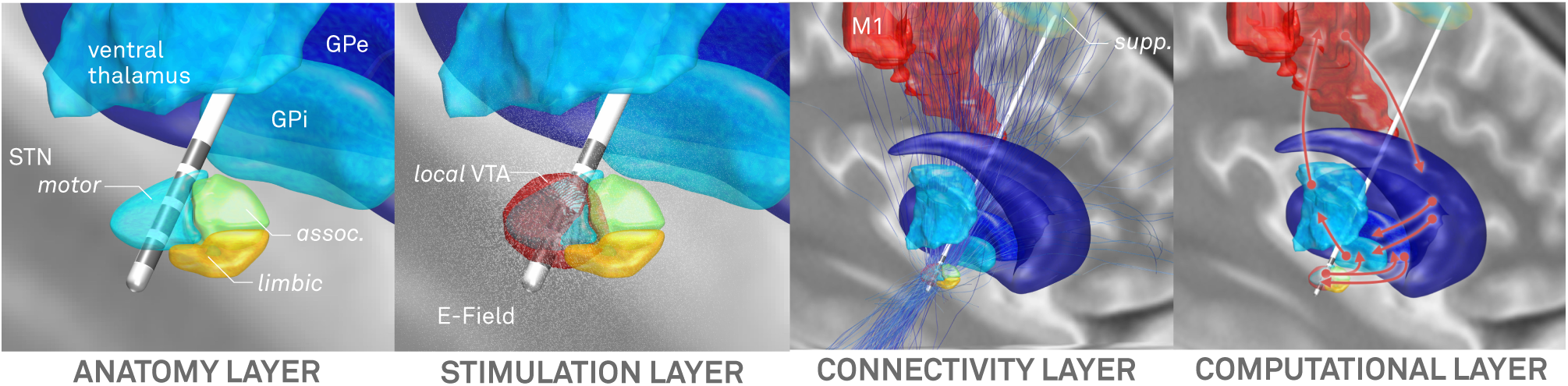
Four layers in a DBS imaging pipeline that may need continuous refinement as technology and methodology advance.

The second layer deals with modeling the local stimulation effects which are often represented by an E-Field or VTA. The anatomy layer directly informs these computations given distinct and even anisotropic conductivity values present in gray or white matter (Butson et al., 2006; Horn et al., 2017c). As mentioned above, to this end, other groups have created much more elaborate models over the last twenty years. Among others, pioneering work by the McIntyre, Butson, Grill, van Rienen and WÅrdell groups should be mentioned (e.g. Åström et al., 2014; 2009; Butson et al., 2006; Butson and McIntyre, 2008; Chaturvedi et al., 2013; Gunalan et al., 2017; Schmidt et al., 2013; Schmidt and van Rienen, 2012). A practical disadvantage of these models is that they require manual interventions at multiple stages and use of a multitude of software applications (some of which are expensive commercial solutions; e.g. see Gunalan et al., 2017). On the other end of the spectrum, even simpler models exist that were successfully employed in clinical context (Dembek et al., 2017; Kuncel et al., 2008; Lauro et al., 2015; Mädler and Coenen, 2012; Vanegas Arroyave et al., 2016). Still, while it remains to be shown that more clinical variance may be explained when applying more sophisticated models, the stimulation layer is definitely one where Lead-DBS has yet much room for improvement.

The third layer deals with the transition from a local VTA to a global volume of modulation by applying brain connectivity. Using tractography or functional connectivity to estimate which other brain areas could potentially be modulated by DBS is a powerful technique that was already used to predict clinical outcome in PD patients (Horn et al., 2017c). However, a big challenge is that both methods are highly indirect. As recently demonstrated, tractography results are dominated by false positive connections (Maier-Hein et al., 2017). On the other hand, resting-state functional MRI is only able to give rough statistical dependencies between an indirect measure of brain activity that operates on a very slow temporal scale. Thus, the conclusions drawn from these measures need careful interpretation and benefit from validation via anatomical or electrophysiological work. For instance, the use of combined LFP-MEG recordings (Litvak et al., 2011; Neumann et al., 2015; Oswal et al., 2016) may validate fMRI findings and vice versa, while animal, tracer or gross-dissection studies may be used to interpret tractography results (e.g. Forel, 1877; Iwahori, 1978; T. Kita and H. Kita, 2012; Marburg, 1904). With these limitations in mind, it should be mentioned that the two main tractography algorithms included in Lead-DBS were each best performers in large open competitions (Fillard et al., 2011; Maier-Hein et al., 2017) and a specific advantage of the GQI method in clinical context lies in its low false positive score (Maier-Hein et al., 2017).

Finally, a fourth layer could be seen as modeling dynamics or connectivity changes induced by DBS. This layer is not touched upon here, but computational modeling based on empirical data seems the only way to investigate how brain activity and connectivity responds to stimulation of a specific target. Already, Lead-DBS was used in such basal ganglia modeling studies (Schroll et al., 2015) and an aim of future versions is to incorporate or interface with modeling software – steadily working toward a “virtual patient” model that facilitates a better understanding of DBS.

In conclusion, we present an updated, advanced and integrative platform for DBS imaging research that is openly available with the aim to further elucidate the mechanisms of DBS and improve therapeutic outcome for DBS patients worldwide.

## Acknowledgements

This study was supported by Deutsche Forschungsgesellschaft (grants KFO 247, SPP 2041) to AAK, Stiftung Charité, Berlin Institute of Health and Prof. Klaus Thiemann Foundation to AH. TMH has received research support from the American Brain Foundation / American Academy of Neurology and NINDS grant K23NS099380. QF was supported by NIH R01-GM114365 (from NIGMS) and R01-CA204443 (from NCI). TP was supported by the Victorian Government’s Operational Infrastructure Support Programme, the Colonial Foundation and the St. Vincent’s Hospital Melbourne Research Endowment Fund. JV was supported by the National Science Foundation (NSF): US IGNITE - 10037840. NL, AK, CB, SE, AT, AH, TP, MR, HS, RO, CR, FCY, QF, TMH and JV have nothing to disclose. AAK reports personal fees and non-financial support from Medtronic, personal fees from Boston Scientific and personal fees from St. Jude Medical outside the submitted work. AH and TAD received speaker honoraria from Medtronic outside the submitted work. WJN received travel grants from Medtronic and St. Jude medical outside of the submitted work. Data collection and sharing for this project was provided by the Human Connectome Project (HCP; Principal Investigators: Bruce Rosen, M.D., Ph.D., Arthur W. Toga, Ph.D., Van J. Weeden, MD). HCP funding was provided by the National Institute of Dental and Craniofacial Research (NIDCR), the National Institute of Mental Health (NIMH), and the National Institute of Neurological Disorders and Stroke (NINDS). HCP data are disseminated by the Laboratory of Neuro Imaging at the University of Southern California. The HCP project (Principal Investigators: Bruce Rosen, M.D., Ph.D., Martinos Center at Massachusetts General Hospital; Arthur W. Toga, Ph.D., University of Southern California, Van J. Weeden, MD, Martinos Center at Massachusetts General Hospital) is supported by the National Institute of Dental and Craniofacial Research (NIDCR), the National Institute of Mental Health (NIMH) and the National Institute of Neurological Disorders and Stroke (NINDS). HCP is the result of efforts of co-investigators from the University of Southern California, Martinos Center for Biomedical Imaging at Massachusetts General Hospital (MGH), Washington University, and the University of Minnesota. Data used in the preparation of this article were obtained from the Parkinson’s Progression Markers Initiative (PPMI) database (www.ppmi-info.org/data). For up-to-date information on the study, visit www.ppmi-info.org. PPMI - a public-private partnership - is funded by the Michael J. Fox Foundation for Parkinson’s Research and funding partners, see www.ppmi-info.org/fundingpartners.

## Supplementary Material

### Imaging parameters of the 3T example patient

Presurgical imaging was performed on a 3T MRI system (Skyra Magnetom, Siemens, Erlangen, Germany). The patient was not sedated for imaging. For better visualization of vascular structures, a standard dose of 0.1 mmol/kg gadobutrol was administered intravenously at the start of the procedure. The MRI protocol consisted of the following sequences: head scout (14 s), sagittal 3D T1-weighted gradient-echo sequence (MPRAGE; TR/TE/TI 2300/2.32/900 ms, voxel size 0.9 × 0.9 × 0.9 mm^3^, FOV 240 × 240 mm^2^; 5 min. 21 s), axial T2 TSE (TR/TE 13320/101 ms, voxel size 0.7 × 0.7 × 2.0 mm^3^, FOV 250 × 250 mm^2^; 3 min. 48 s), sagittal Fast Grey Matter Acquisition T1 Inversion Recovery (FGATIR; TR/TE/TI 1500/3.56/559 ms, voxel size 0.8 × 0.8 × 1.0 mm^3^, FOV 240 × 240 mm^2^; 6 min. 17 s), axial proton-weighted TSE (TR/TE 3600/11 ms, voxel size 1.0 × 1.0 × 2.0 mm^3^, FOV 250 × 250 mm^2^; 3 min. 34 s), and dMRI (EPI, TR/TE 7500/95 ms, voxel size 2.0 × 2.0 × 2.0 mm^3^, FOV 250 × 250 mm^2^, 20 diffusion-encoding directions with b-value 1000 s/mm^2^ and one non-diffusion-weighted b_0_-volume; 5 min. 47 s). Overall acquisition time was 25 min. A routine head CT was done the day after electrode implantation using an 80-multislice Toshiba Aquilion PRIME (Canon Medical Systems, Otawara, Japan). Four sequential blocks, each with a length of 37.75 mm, were acquired (280 mA, 120 kV, rotation time 1 s, matrix 512 x 512 mm^2^, CTDI_vol_ 55.79 mGy), resulting in a slice thickness of 0.5 mm.

### Imaging parameters of the retrospective 1.5T cohort

All patients underwent pre-operative MR-imaging on a 1.5 T scanner (NT Intera; Philips Medical Systems, Best, The Netherlands) using a T2-weighted fast spin-echo (FSE) with the following parameters: TR = 3500 ms, TE = 138 ms, echo-train length: 8, excitations: 3, flip angle: 90°, section thickness: 2 mm, section gap: 0.2 mm, FOV: 260 mm (in-plane resolution 0.51 × 0.51 mm), matrix size: 384 interpolated to 512, total acquisition time, 10 min and 41 s.

Postoperative MR-imaging was performed in 45 patients. DBS patients are subject to a limitation of the specific absorption rate (SAR, b0.1 W/kg), which has been specified by the manufacturer of the electrodes. Within 5 days after implantation of the electrodes, MR-imaging was performed on the same scanner using a T2-weighted fast spin-echo (FSE) sequence in low SAR mode with the same parameters as used pre-operatively. Philips software Version 11.1 level 4 was used. MR sections in the axial and coronal planes were obtained and processed in this study. In the following, “axial” and “coronal” volumes refer to acquisitions with voxel sizes of 0.51 × 0.51 mm in the axial or coronal planes respectively, each with a 2 mm slice thickness. Postoperative CT was conducted in 6 patients (8 of whom also had postoperative MRI). Here, high-resolution images were acquired on a LightSpeed16 (GE Medical System, Milwaukee, Wisconsin) Slice CT with a spatial resolution of 0.49 × 0.49 × 0.67 mm^3^. Images were acquired in axial (i.e. sequential/incremental) order at 140 kV and automated mA setting. Noise index was 7.0. A large SFOV with 50 cm diameter was used.

### Methodological details for normalization strategies as implemented in Lead-DBS

#### SPM Unified Segmentation

In Lead-DBS, the Unified Segmentation approach does not fit warps to the standard tissue probability maps (TPM) supplied with SPM software which are based on a population differing from the original MNI-152 population (IXI-Dataset; http://brain-development.org/ixi-dataset/). Instead, the multispectral (T1-, T2-, PD-weighted) MNI-152 / ICBM 2009b Nonlinear Asymmetric template series were preprocessed using the Unified Segmentation approach, generating a new set of templates. Moreover, this was done using refined tissue priors that include subcortical structures and have been proposed for use in the DBS context (Lorio et al., 2016). This procedure ensures direct compatibility with the 2009b space (Fonov et al., 2009) and the use of these adapted TPM yielded slightly better segmentations of STN and GPi in a recent comparison (Ewert et al., 2018a). Of note, all implementations of SPM methods (Unified Segmentation, DARTEL & SHOOT, see below) within Lead-DBS by default operate on all preoperative acquisitions simultaneously (multispectral normalization).

#### SPM DARTEL & SHOOT

SPM DARTEL/SHOOT create population-specific group average templates based on the data available (Ashburner, 2007; Ashburner and Friston, 2011). As these are different to MNI space, we adapted the pipeline to register to single subjects directly to MNI space derived from the multispectral 2009b data described above.

#### Advanced Normalization Tools (ANTs)

ANTs is another non-linear pipeline that has been extensively optimized for LEAD-DBS 2.0 (Ewert et al., 2018a). The current default implementation is a multispectral warp that combines a rigid, an affine and two nonlinear symmetric image normalization (SyN) stages. While the first SyN stage includes the whole brain, the second is limited to a subcortical region mask defined in (Schönecker et al., 2009). The implementation detects and properly handles slabs, which are often used in clinical context. For instance, many centers acquire whole-brain 3D gradient-echo T1-weighted series and a T2-weighted slab of the subcortex for stereotactic planning (Horn et al., 2017c). If slabs are detected, Lead-DBS automatically adds a third intermediate SyN stage that focuses on the area covered by all acquisitions (whereas the linear stages and first SyN stage only takes the whole-brain acquisition(s) into account).

#### MAGeT Brain-like Segmentation

The MAGeT Brain approach (https://github.com/CobraLab/MAGeTbrain; Chakravarty et al., 2012) was developed for segmentation of deep nuclei and computes multiple warps from atlas space via multiple “templates” to the subject of question. Here, templates refer to additional brains (e.g. high resolution MRIs acquired on specialized hardware). Thus, the deformation from atlas to subject space takes “detours” via multiple different brains and, as a result, multiple solutions for the atlas based segmentation are attained. These solutions are then aggregated (e.g. by simple averaging or majority voting). The intuitive understanding of the approach is that it increases robustness by introducing anatomical variance and the original authors showed this increased precision empirically in multiple studies (Chakravarty et al., 2012; Pipitone et al., 2014).

In Lead-DBS, this concept is adopted and used for both segmentation and normalization. In a first step, key structures defined by the DISTAL atlas in 2009b space (STN, GPi, GPe, RN) are segmented in native space using an adapted MAGeT Brain approach implemented in Lead-DBS using ANTs. In a second step, a SyN-based ANTs-registration from native to template space is computed (see above) that additionally includes the segmented labels as further “spectra” in the warp. For example, if preoperative T1- and T2-weighted acquisitions are present, a six-fold normalization is computed (including T1-patient to T1-MNI, T2-patient to T2-MNI, auto-segmented patient STN to DISTAL atlas STN, patient GPi to DISTAL GPi, etc.). This approach may add additional precision at and around key interest regions in DBS but has not been empirically tested or compared to conventional approaches beyond the original MAGeT publications (Chakravarty et al., 2012; Pipitone et al., 2014).

#### MAGeT Brain-like Normalization

This further adaptation of the MAGeT Brain approach reverses the idea and directly works on the deformation fields solved during normalization. Specifically, multiple deformation fields warping from subject to template are created, each taking a detour via a peer brain (“template” in the MAGeT-Brain nomenclature). Resulting deformation fields are then aggregated by calculating their voxel-wise robust average. A final deformation field that warps from subject to template space is created that can be applied to warp electrodes into template space. Of note, both MAGeT-inspired methods require the use of “peer-subjects” to include into the analysis. To demonstrate this in the present patient example, 21 age-matched peers from the IXI-dataset (http://brain-development.org/ixi-dataset/) were automatically assigned by Lead-DBS as previously described in similar context (Horn et al., 2017a).

### Methodological details for volume of tissue activated generation

#### Mesh Generation

A cylindrical four-compartment model of the area surrounding the electrode and the lead itself is constructed based on the conducting and insulating parts of the electrodes, gray matter, and white matter. This model is represented by a tetrahedral mesh that is generated using the iso2mesh toolbox (http://iso2mesh.sourceforge.net/) in combination with other meshing tools, including Tetgen (http://wias-berlin.de/software/tetgen/) for the Delaunay tetrahedralization step and Cork (https://github.com/gilbo/cork) for merging electrode and brain surface meshes. We note here that all involved tools are included within Lead-DBS and some interfaces of iso2mesh were slightly modified. The output mesh is truncated by a cylindrical boundary with a radius of 20 mm; a larger radius is automatically used when higher stimulation amplitudes are applied. All regions within the cylinder not covered by either gray matter or the electrode are modeled as isotropic white matter. Gray matter portions are defined by the atlas chosen by the user (table 5). This gray matter model can be either informed by an atlas available in template space, the same atlas that was automatically transferred to native space or a manual segmentation of structures of interest manually performed by the user.

Finally, the conducting and insulating regions are specified by the electrode models supplied with Lead-DBS. A wide range of electrodes from five manufacturers are readily implemented in Lead-DBS (table 7) and it is straight forward to implement custom models. Surface meshes from electrode and atlas as well as the bounding cylinder are resolved to a joint surface mesh using a modified version of Cork as implemented in iso2mesh. This joint mesh is then fed into Tetgen to create a tetrahedral mesh. Tetrahedra within the regions of gray matter, white matter, metal or insulating parts are identified and tagged using a post-processing code implemented in Lead-DBS.

#### Electric Field Modeling using Finite Element Method (FEM)

Conductivity at each tetrahedron is estimated using the FieldTrip / SimBio pipeline (Oostenveld et al., 2010; Vorwerk et al., 2014; 2018; https://www.mrt.uni-jena.de/simbio/; http://www.fieldtriptoolbox.org/). Conductivity values for gray and white matter can be entered by the user while two presets are available. The first consists of conductivity values commonly used in the field (0.33 and 0.14 S/mm for gray and white matter, respectively). Similar values were used in modeling EEG data (e.g. Buzsáki, 2006; Vorwerk et al., 2014) and in most previous DBS models (e.g. Åström et al., 2014; McIntyre and Grill, 2002). However, these conductivity values are not corrected for stimulation frequency and drop drastically at a typical stimulation frequency of 130 Hz (Hasgall et al., 2015). Thus, a second set of conductivity presets adjusted for stimulation frequency is available (0.0915 and 0.059 S/mm).

Both constant-voltage and constant-current settings can be modeled. The voltage or amplitude applied to the active contacts is introduced as a boundary condition. In case of monopolar stimulation, the surface of the bounding cylinder serves as the anode. Subsequently, the gradient of the potential distribution is calculated by derivation of the FEM solution. Due to the first order FEM approach that is used, the resulting gradient is piecewise continuous. The binary VTA is obtained by thresholding the resulting electric field strength at a user-specified value. In the past, a value of 0.2 V/mm has frequently been used in this context (Åström et al., 2014; 2009; Horn et al., 2017c; Vasques et al., 2009). In Lead-DBS, this value serves as the “general heuristic” default value. Furthermore, based on prior literature, heuristics for STN (0.19 V/mm; Åström et al., 2014; Mädler and Coenen, 2012), GPi (0.2 V/mm; Hemm et al., 2005; Vasques et al., 2009) and Vim (0.165 V/mm; Åström et al., 2014; Kuncel et al., 2008) can be used. Finally, specific activation thresholds that depend on axon diameters (D = 2-5 μm) and stimulation pulse-widths (t = 30, 60, 90, 120 μs) are available as they were estimated in (Åström et al., 2014).

**Figure S1:**
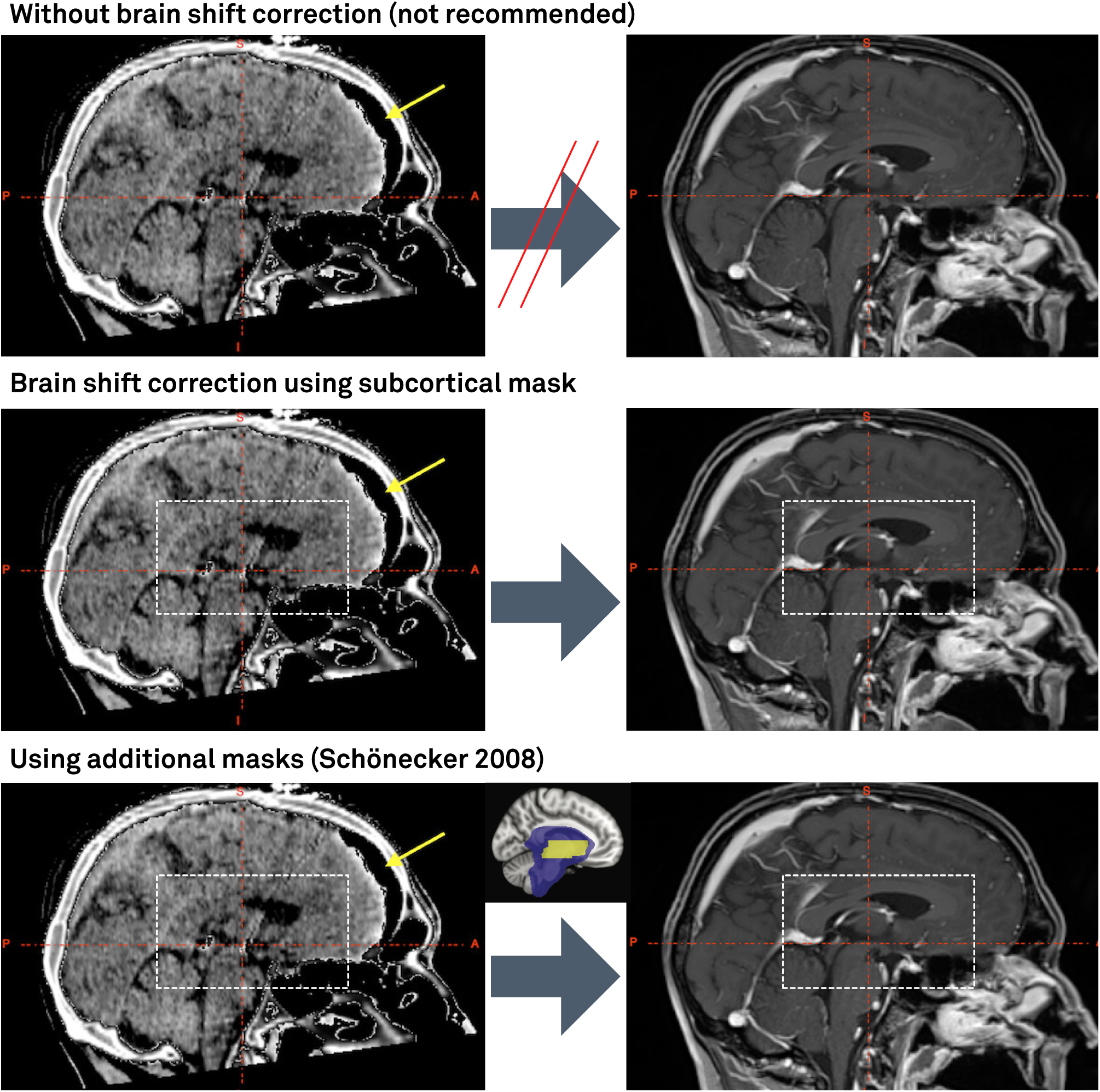
Brain shift correction approach. The top panel shows a whole-brain affine registration aligning a postoperative CT with frontal pneumocephalus (left) to a preoperative T1-weighted MRI (post gadolinium). Based on the pneumocephalus (yellow arrow), a nonlinear error results that is largest in frontal regions (at the site of the pneumocephalus) that may still be substantial in the regions of interest of DBS (figure 4). In the mid row, an additional refinement transform between the subcortical area delineated by the white dashed box is computed. Finally, in the third row, one or two additional refinement transforms between subcortical areas of interest are computed based on masks defined in (Schönecker et al., 2009). This approach gradually shifts the registration area away from the area nonlinearly distorted by pneumocephalus toward the subcortical regions of interest.

## List of abbreviations

ANTs: Advanced Normalization Tools, see http://stnava.github.io/ANTs/
BSpline-SyN: ANTs normalization method; Explicit B-spline regularization in symmetric diffeomorphic image registration
CORE: Algorithm for “reconstruction of electrode contact positions” defined in Horn & Kühn 2014
DARTEL: SPM normalization method; A fast diffeomorphic image registration algorithm.
DiODe: Directional Orientation Detection (Sitz et al. 2017)
dMRI: Diffusion-weighted MRI – in the context of diffusion-imaging based tractography.
FA: Fractional Anisotropy
FGATIR: Fast Grey Matter Acquisition T1 Inversion Recovery
FLIRT: FMRIB’s Linear Image Registration Tool
FNIRT: FSL normalization method; FMRIB’s Nonlinear Image Registration Tool
FSL: FMRIB Software Library, see https://fsl.fmrib.ox.ac.uk/
GNU: Recursive acronym for “GNU’s Not Unix.”; GNU GPL is a popular open license supported by the Free Software Foundation (see http://www.gnu.org). The Free Software Foundation (FSF) is a nonprofit with a worldwide mission to promote computer user freedom.
Gpi: internal segment of the globus pallidus
Gpe: external segment of the globus pallidus
GQI: Generalized q-sampling imaging, dMRI processing method implemented in DSI studio
HCP: Human Connectome Project
LEDD: Levodopa Equivalent Daily Dosage
MAGeT Brain: Multiple Automatically Generated Templates brain segmentation algorithm, see https://github.com/CobraLab/MAGeTbrain
PaCER: Precise and Convenient Electrode Reconstruction for DBS, see https://adhusch.github.io/PaCER/
PCA: Principal Component Analysis
PPMI: Parkinson‘s Disease Progression Marker Initiative
PPN: Pedunculopontine nucleus
RN: red nucleus
ROI: Region of Interest
RSME: Root-mean-square error
QSM: Quantitative Susceptibility Mapping
rs-fMRI: Resting-state functional MRI
SHOOT: SPM normalization method; Diffeomorphic registration using geodesic shooting and Gauss–Newton optimisation
SPM: Statistic Parametric Mapping, see http://www.fil.ion.ucl.ac.uk/spm/software/spm12/
STN: subthalamic nucleus
SyN: ANTs normalization method; Symmetric diffeomorphic image registration / symmetric image normalization
TRAC: Algorithm for “trajectory reconstruction” defined in Horn & Kühn 2014
UPDRS: Unified Parkinson‘s Disease Rating Scale
VTA: Volume of Tissue Activated

